# Sister-chromatid-sensitive Hi-C reveals the conformation of replicated human chromosomes

**DOI:** 10.1101/2020.03.10.978148

**Authors:** Michael Mitter, Catherina Gasser, Zsuzsanna Takacs, Christoph C. H. Langer, Wen Tang, Gregor Jessberger, Charlie T. Beales, Eva Neuner, Stefan L. Ameres, Jan-Michael Peters, Anton Goloborodko, Ronald Micura, Daniel W. Gerlich

## Abstract

The three-dimensional organization of the genome supports regulated gene expression, recombination, DNA repair, and chromosome segregation during mitosis. Chromosome conformation capture (Hi-C)^1–3^ has revealed a complex genomic landscape of internal chromosome structures in vertebrate cells^4–11^ yet how sister chromatids topologically interact in replicated chromosomes has remained elusive due to their identical sequences. Here, we present sister-chromatid-sensitive Hi-C (scsHi-C) based on nascent DNA labeling with 4-thio-thymidine. Genome-wide conformation maps of human chromosomes revealed that sister chromatid pairs interact most frequently at the boundaries of topologically associating domains (TADs). Continuous loading of a dynamic cohesin pool separates sister-chromatid pairs inside TADs and is required to focus sister chromatid contacts at TAD boundaries. We identified a subset of TADs that are overall highly paired, characterized by facultative heterochromatin, as well as insulated topological domains that form separately within individual sister chromatids. The rich pattern of sister chromatid topologies and our scsHi-C technology will make it possible to dissect how physical interactions between identical DNA molecules contribute to DNA repair, gene expression, chromosome segregation, and potentially other biological processes.

---

Expression, maintenance, and inheritance of genetic information relies on highly regulated topological interactions within and between huge chromosomal DNA molecules. For instance, intramolecular contacts between promoters and distant enhancers activate gene transcription^12^, whereas intermolecular contacts between homologous DNA sequences of replicated sister chromatids enable error-free DNA damage repair^13^. During cell cycle progression, conformational changes within and between the replicated sister chromatids shape mechanical bodies that can be segregated by the mitotic spindle^7–11,14^. In vertebrates, intramolecular DNA loops are dynamically formed by the Structural Maintenance of Chromosomes complex cohesin^15–21^ within boundaries established by the CCCTC-binding factor (CTCF), thereby structuring chromosomes into topologically associating domains (TADs)^4–6^. This organization contributes to transcriptional control^12,22^, and misregulated chromosome conformations have been associated with developmental disorders^23,24^ and cancer^25,26^. A distinct pool of cohesin links sister chromatids^27^ topologically to enable homology-directed DNA repair^28,29^ and chromosome segregation in subsequent mitosis^30–32^. However, it is not known how cohesive linkages distribute on the genome to support these important functions, and how they are coordinated with dynamic loop formation and TADs.

The complex organization of vertebrate genomes has been revealed by chromosome conformation capture technology (Hi-C)^1–3^, which maps DNA contacts genome-wide. The development of Hi-C led to the discovery of TADs^4–6^ and revealed how they are dynamically remodeled during the cell cycle^7–11^. It also allowed the elucidation of how cohesin regulates dynamic loop formation^15–19^, and it has been widely used to study the functional implications of chromosome conformation in various biological contexts. However, Hi-C technology cannot currently be used to explore topological interactions between the sister chromatids of replicated chromosomes, as the identical DNA sequences in replicated chromosomes make it impossible to distinguish between intra-molecular and inter-molecular contacts. To overcome this limitation, we have developed sister-chromatid-sensitive Hi-C (scsHi-C) for genome-wide conformation analysis of replicated human chromosomes.

## Sister-chromatid-sensitive Hi-C

To distinguish between cis and trans sister chromatid contacts, it would be necessary to introduce a sister-chromatid-specific label. We reasoned that this could be achieved by culturing cells for one round of DNA replication in the presence of a DNA nucleotide analogue to label the Watson strand on one sister chromatid and the Crick strand on the other (Fig. 1a). If the nucleotide analogue can be detected by DNA sequencing, then standard Hi-C procedures^6,33^ could be used to categorize chromatid contacts as either cis, which would be labelled on the same strand, or trans, which would be labelled on different strands (Fig. 1b).

**Figure 1.**
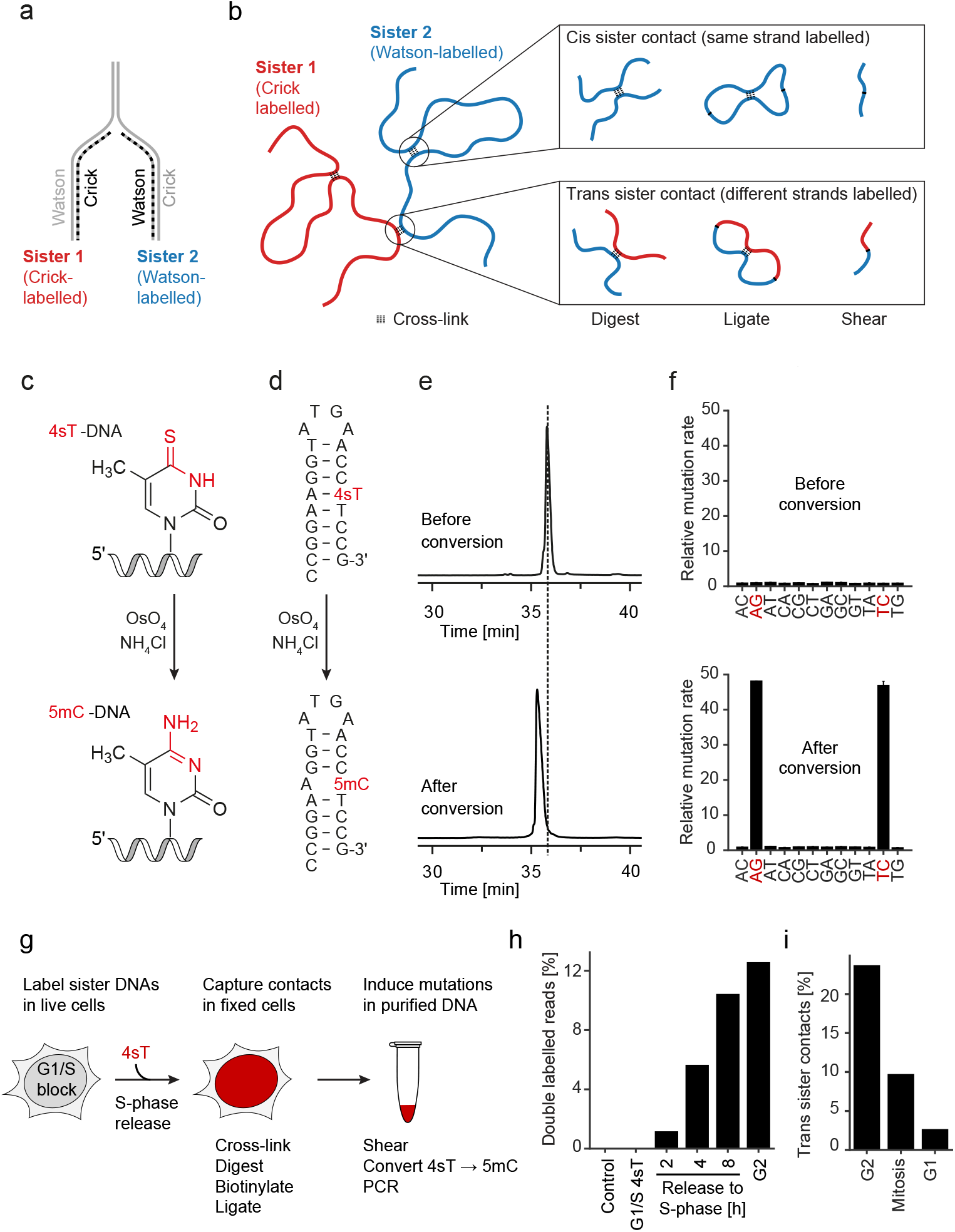
scsHi-C methodology based on nascent DNA labeling in live cells. (a) Sister chromatid-specific labeling using synthetic nucleotides. During DNA replication, a synthetic nucleotide analogue incorporates into different strands (Watson or Crick) within each sister chromatid. Labelled DNA, dashed line; unlabelled DNA, solid line. (b) Strategy to distinguish cis from trans sister contacts in a Hi-C experiment based on 4sT-mediated DNA labeling. After progression through S-Phase in the presence of 4sT, each sister chromatid contains one labelled DNA strand of opposing strandedness (see panel a). Chromatin is crosslinked in cells and Hi-C samples are prepared using standard procedures, followed by chemical conversion to induce 4sT signature mutations and Illumina-based sequencing. Half-reads are classified as labelled if at least two signature mutations are present. If a ligation junction contains two labelled half-reads that map to the same strand, it is classified as cis sister contact; if it contains two labelled halves that map to opposing strands, it is classified as a trans sister contact (see Extended Data Fig. 3c for details). (c) Conversion of 4-thio-thymidine (4sT) to the point-mutation inducing 5-methyl-Cytosine (5mC) by treating DNA with OsO_4_ and NH_4_Cl at elevated temperatures. Functional groups that are changed in the course of the reaction are highlighted in red. (d) Synthetic hairpin-oligonucleotide used to probe 4sT conversion by OsO_4_. The theorized reaction educts and products are highlighted in red. (e) High performance liquid chromatography (HPLC) trace at 260 nm of the oligos depicted in (d) before and after the conversion by OsO_4_/NH_4_Cl. The peak position of the oligo before conversion is indicated by a dashed line. (f) Point-mutation rates of genomic DNA from HeLa cells grown in medium containing 4sT relative to control DNA from cells grown in the absence of 4sT, before and after OsO_4_/NH_4_Cl-mediated conversion. Bar graphs indicate the mean and standard error of three independent experiments. (g) Experimental procedure for differential labeling of sister chromatids using 4sT. See Extended Data Fig. 2c for more details. (h) Quantification Hi-C reads that are labelled on both sides for contact sister-specificity classification, as a percentage of all reads. Cells were synchronized to the G1/S boundary and released into S-Phase in the presence of 4sT for the indicated times. The G2 sample was arrested using RO3306; the control sample refers to unlabelled DNA. Bars show the mean of two biological replicates. (i) Percentage of trans sister contacts based on all double-labelled reads that exhibit a genomic separation larger than 10kb. Cells were released from G1/S block into medium containing 4sT and then arrested in G2 using RO3306, in mitosis using nocodazole, or the following G1 using thymidine. Bars show mean of two biological replicates.

To establish a sister-chromatid-specific DNA label, we considered 4-thio-thymidine (4sT), because its RNA-analogue 4-thio-uridine is used to label nascent RNA without compromising cell viability^34,35^. ThioUridine-to-Cytidine-sequencing (TUC-seq)^35–37^ employs 4-thio-uridine labeling combined with post-extraction OsO_4_ / NH_4_Cl conversion chemistry to generate distinct point mutations. If this chemistry could be adapted to convert 4sT into 5-methyl-cytosine (5mC) after genomic DNA purification (Fig. 1c), then 4sT-labelled DNA would also generate signature mutations that could be detected by high-throughput sequencing. First, we tested whether 4sT within synthetic DNA oligonucleotides can be converted into 5mC by OsO_4_ / NH_4_Cl chemistry and found that, after 3 hours, virtually all 4sT was converted into 5mC (Fig. 1d, e; Extended Data Fig. 1a, b). To assess toxicity, we cultured HeLa cells in medium containing 4sT and found that 4sT did not activate a DNA damage response and did not compromise cell viability up to 6 mM 4sT (Extended Data Fig. 1c, d). Furthermore, 2 mM 4sT only slightly prolonged S-phase, and almost all cells progressed through mitosis to the following G1 phase (Extended Data Fig. 1e). Thus, 4sT fulfils key requirements for sister-chromatid-specific labelling of genomic DNA in live cells.

Next, we assessed how efficiently 4sT labels genomic DNA. Cells were grown in the presence or absence of 2 mM 4sT for 5 days to establish fully labelled and unlabelled conditions, respectively; genomic DNA was purified, treated with OsO_4_ / NH_4_Cl, amplified and sequenced. Conversion of 4sT to 5mC should yield reads with signature A-to-G and T-to-C point mutations depending on whether the forward or reverse strand of a PCR amplicon is sequenced. Indeed, these signature mutations were elevated almost 50-fold in DNA isolated from 4sT-treated cells compared to controls, whereas other point mutations were not affected (Fig. 1f). The overall frequency of signature mutations was 2.5%, in agreement with mass spectrometry-based detection of 4sT in genomic DNA (Extended Data Fig. 2a, b). Importantly, 4sT-labelled DNA that was not chemically converted had a similar point mutation distribution to that of unlabelled DNA (Fig. 1f). Thus, chemical conversion of 4sT-labelled genomic DNA produces a strong and specific mutation signature that can be detected by high-throughput sequencing.

To implement an scsHi-C procedure based on 4sT labeling (Fig. 1g), we synchronized HeLa cells to the G1/S boundary, released them into medium containing 4sT for one S-Phase, and arrested them in the subsequent G2 phase using the Cdk1 inhibitor RO3306^38^ (Extended Data Fig. 2c, d). Chromatin was cross-linked, digested, tagged with biotin, ligated and purified, as per standard Hi-C procedures adapted from Mumbach et al^33^; 4sT was converted to 5mC and DNA libraries were prepared for high-throughput sequencing. To assess whether 4sT labeling impairs Hi-C analysis, we constructed Hi-C maps from all contacts and found that they closely resembled those from similarly processed cells grown without 4sT treatment (Extended Data Fig. 2e, f). Thus, 4sT labelling and chemical conversion does not perturb genome conformation.

To construct sister-chromatid-resolved contact maps, we can use only Hi-C contacts that contain 4sT-specific point mutations on both sides of the contact (“double-labelled reads”), since at 2.5% incorporation density of 4sT into genomic DNA, the absence of signature mutations does not allow reliable assignment to labelled or unlabelled strands of a sister chromatid (Extended Data Fig. 3a). To assess how many signature mutations are required to confidently detect double-labelled reads, we analyzed Hi-C libraries from cells grown in the absence of 4sT. When considering reads that contained at least two signature mutations, less than 0.2% were classified as “double labelled”, indicating very low rates of misclassification (Extended Data Fig. 3b). When cells were released into S-Phase in the presence of 4sT, the percentage of double-labelled reads increased to 12% in the subsequent G2 phase (Fig. 1h). Thus, double-labelled reads are detected with a very low false positive rate and at sufficient yield to construct Hi-C-maps.

Reads were then classified as cis sister contacts if the labels mapped to the same chromosomal DNA strand on both sides, and as trans sister contacts if the labels mapped to different chromosomal DNA strands on each side (Fig. 1b; Extended Data Fig. 3c). Based on a statistical procedure from Erceg et al^37^, we estimated that wrongly assigned trans sister contacts are below 2 % (Extended Data Fig. 3d,e). Thus, scsHi-C enables accurate discrimination between cis- and trans sister contacts.

To assess whether scsHi-C can detect global resolution and segregation of sister chromatids, we analyzed cells progressing from G2 through mitosis to the following G1 (Extended Data Fig. 2c, d). Sister chromatids resolve during mitotic entry, which should reduce trans sister contacts. In the following G1, each daughter cell inherits only one labelled sister chromatid per homologous chromosome and trans sister contacts should not be present. scsHi-C analysis indeed showed that trans sister contacts dropped from 23.7 % in G2 to 9.8 % in prometaphase and 2.7 % in the following G1 (Fig. 1i), indicating that scsHi-C enables genome-wide analysis of sister chromatid interactions in human cells.

## Conformation of replicated chromosomes

In interphase nuclei, replicated chromosomes must be organized in a way that allows efficient sister chromatid interactions for homology-directed DNA repair, whereas during mitosis, sister chromatids should be largely disentangled so that they can be easily moved apart by the mitotic spindle. To determine where sister chromatids contact each other during interphase and to measure the extent of sister chromatid resolution during mitosis, we constructed genome-wide scsHi-C maps of cells synchronized to G2 or mitotic prometaphase (Fig. 2a, b; Extended Data Fig. 2c, d; Table S1, 2). G2 maps were constructed based on 1.7 billion Hi-C reads, which yielded 195 million unique sister-chromatid-specific contacts of 11 highly reproducible replicates (Extended Data Fig. 4a, b; Table S1). In low-magnification views, trans sister contacts were highly enriched along the diagonal (Fig. 2a), indicating that sister chromatids were overall aligned, thereby favoring interactions between homologous genomic regions during G2. In scsHi-C maps of prometaphase cells, trans sister contacts were very scarce and not substantially enriched along the diagonal (Fig. 2b; Extended Data Figure 4c, d, Table S2), indicating almost complete separation of sister chromatids. Furthermore, cis sister contacts reorganized from a blocked distribution along the diagonal in G2 to a more homogeneous and broader distribution in prometaphase, as previously observed^7,9^. scsHi-C thus reveals globally paired organization of sister chromatids in interphase and almost complete separation in mitosis.

**Figure 2.**
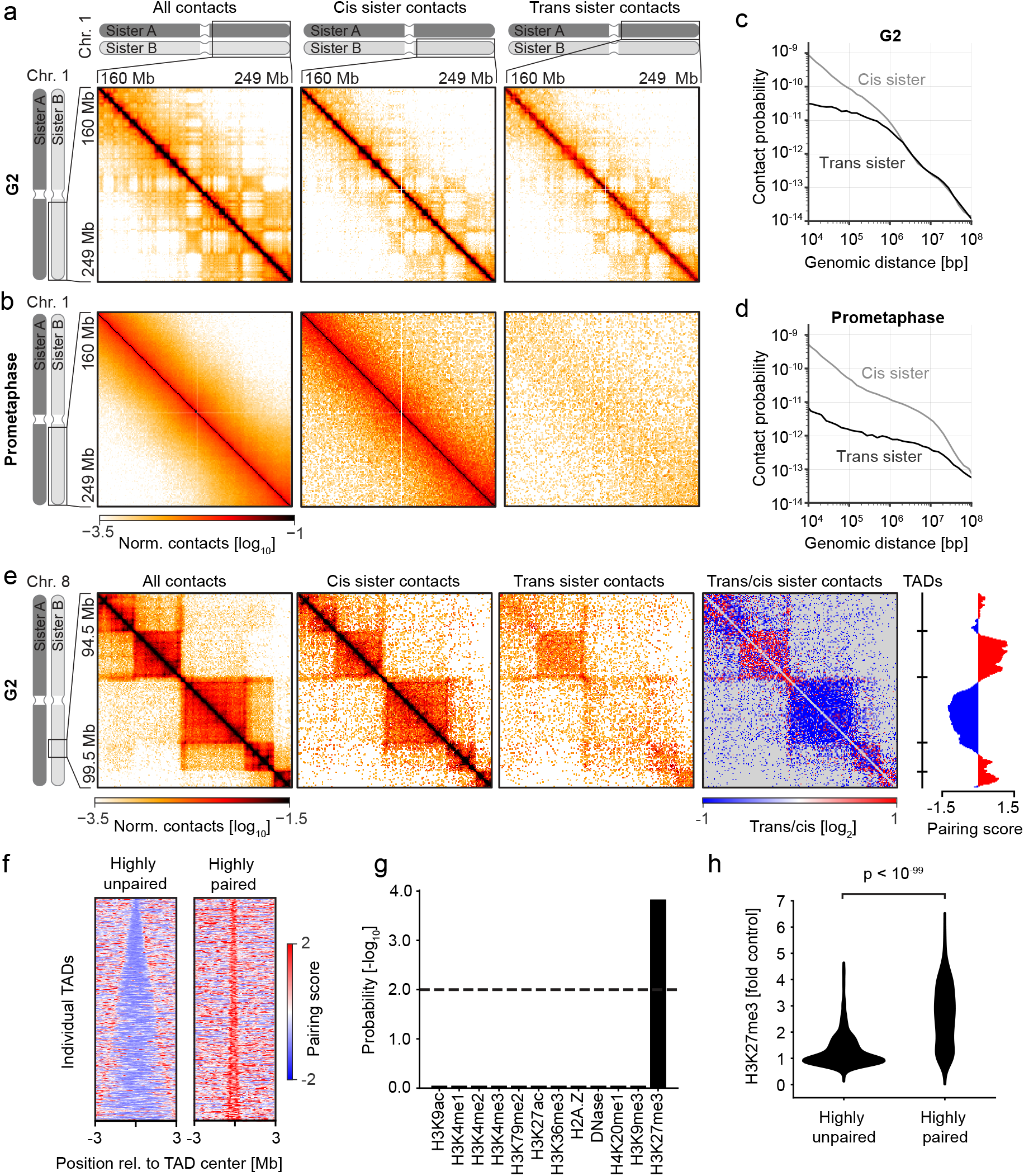
Genome-wide conformation maps of replicated human chromosomes. (a) Hi-C interaction matrices of the long arm of chromosome 1 of all contacts, cis sister-, and trans sister contacts in 11 merged G2 samples. The all-contacts matrix was normalized to the total number of corrected contacts in the region of interest (ROI), whereas cis sister and trans sister contacts were normalized to the total amount of cis sister and trans sister contacts in the ROI. Bin size of the matrix is 500kb. (b) Hi-C interaction matrix of the long arm of chromosome 1 of all, cis sister, and trans sister contacts in two merged prometaphase samples. Contacts were normalized as in (a). Bin size of the matrix is 500kb. (c) Average contact probability over different genomic distances for cis sister and trans sister contacts of the G2 sample shown in (a). (d) Average contact probability as in (c) of the prometaphase sample shown in (b). (e) All-contacts, cis sister and trans sister contacts, as well as the ratio of trans sister observed/expected to cis sister observed/expected of 11 merged G2 samples at a representative region on chromosome 8 is displayed alongside the location of TAD boundaries and the trans sister pairing score within a sliding diamond of 400kb (see Method section for details). Bin size of the matrix is 20kb. (f) Stack-up of trans sister pairing score (see Method section for details) along TADs that are highly paired or highly unpaired (see Extended Data Fig. 5d), sorted by the size of TADs. Shown are windows of 6Mb around the center of the respective TADs. Pairing scores were calculated within a sliding window of 200kb on a Hi-C matrix with 20kb bin size. (g) Visualization of enrichment analysis that was done on TADs that exhibit high pairing (see Fig.S5d) using LOLA ^42^. The panel shows all chromatin modification datasets in the extended LOLA database for HeLa cells with their respective p-value. P-value cut-off (p < 0.01) is displayed as a dashed line. (h) Quantification of H3K27me3 enrichment at highly paired and highly unpaired TADs, displaying the average fold-enrichment of H3K27me3 within the respective intervals. P-value was calculated using a two-sided Mann-Whitney U test.

To investigate sister chromatid organization in more detail, we calculated average contact frequencies over distinct genomic intervals. In G2 cells, cis sister contacts were much more frequent than trans sister contacts over genomic distances up to ~3 Mb (Fig. 2c), indicating extensive local sister separation within the globally paired arrangement. Over larger genomic distances, however, cis and trans sister-contact frequencies were indistinguishable, suggesting an overall intermixed arrangement of sister chromatids in G2 nuclei. In prometaphase cells, cis sister contacts dominated trans sister contacts even at 100 Mb genomic intervals (Fig. 2d), indicating resolution of entire chromosome arms. scsHi-C thus reveals that the locally separated but globally entangled sister chromatids of interphase nuclei convert into almost completely resolved bodies during mitosis.

Given the local separation of sister chromatids in interphase nuclei, we explored how trans sister contacts are coordinated with intra-chromatid conformations. High-magnification scsHi-C maps of G2 chromosomes showed a highly structured pattern of trans sister contacts along the diagonal, which to some extent corresponded to the positions of TADs in cis sister contact maps (Fig. 2e). However, trans and cis sister contact distributions were highly divergent, with some TADs densely filled with trans sister contacts, indicating extensive pairing, whereas others were virtually devoid of trans sister contacts, indicating loose association (Fig. 2e, Extended Data Fig. 5a). Importantly, regions with low frequencies of trans sister contacts detected by scsHi-C correlated well with a high propensity of replicated sister loci to split, as previously observed in the same cell type by fluorescence in situ hybridization and live-cell imaging of dCas9-EGFP^40^ (Extended Data Fig. 5b, c), thus validating our scsHi-C measurements. TADs thus demarcate discrete domains with variable degrees of sister chromatid pairing.

To understand the molecular basis of pairing domains, we annotated TADs in our scsHi-C maps using OnTAD^41^ to quantify a pairing score for each TAD based on average trans sister contact frequency (Fig. 2e; Extended Data Fig. 5d). This revealed hundreds of TADs that were very highly paired, as well as many highly unpaired TADs (Fig. 2f). While the highly paired TADs were apparently smaller in size than the unpaired TADs, we also aimed to identify potential differences in chromatin composition. We therefore correlated the pairing score of individual TADs with different chromatin features using Locus Overlap Analysis (LOLA)^42^. This analysis showed that highly paired TADs were markedly enriched in H3K27me3 (Fig. 2g, h), a mark for polycomb-repressed facultative heterochromatin^43^. The overall degree of sister chromatid pairing within TADs is thus defined by characteristic chromatin modifications.

## Organization of TADs in replicated chromosomes

We next investigated how chromatin fibers fold within individual TADs to comply with dynamic loop formation in the presence of cohesive linkages between sister chromatids. Cis sister contacts were most prominently enriched along the diagonal throughout TADs, indicating high abundance of short-range intra-molecular contacts (Fig. 3a; Extended Data Fig. 6a-c). In contrast, trans sister contacts filled TAD areas without substantial accumulation along the diagonal (Fig. 3a; Extended Data Fig. 6a-c), indicating that sister DNAs are not strictly aligned in a “railroad” configuration^39^ within TADs, despite their globally paired organization. Quantification of contact densities showed that trans sister contacts enriched at many TAD boundaries, whereas cis sister contacts were slightly less abundant (Fig. 3b), indicating that TAD boundaries might be sites where sister chromatids contact most frequently. To address this possibility, we analyzed all TADs annotated in our scsHi-C maps. Aggregated contact probability maps of TADs confirmed trans sister contact enrichment at the diagonal position of TAD boundaries, and less prominently also at corner positions connecting the neighboring TAD boundaries (Fig. 3c). This is consistent with an overall registration of TADs, where a given TAD boundary is in proximity to the same TAD boundary on its sister chromatid, as well as the neighboring TAD boundary of the sister chromatid (Fig. 3d). Analysis of individual TADs by contact density line profiles confirmed that trans sister contacts generally enriched at the boundaries irrespective of the size of TADs (Fig. 3e), on average 2-fold compared to the genome-wide average (Fig. 3f). Thus, sister chromatids are predominantly linked at TAD boundaries, whereas they separate extensively inside TADs.

**Figure 3.**
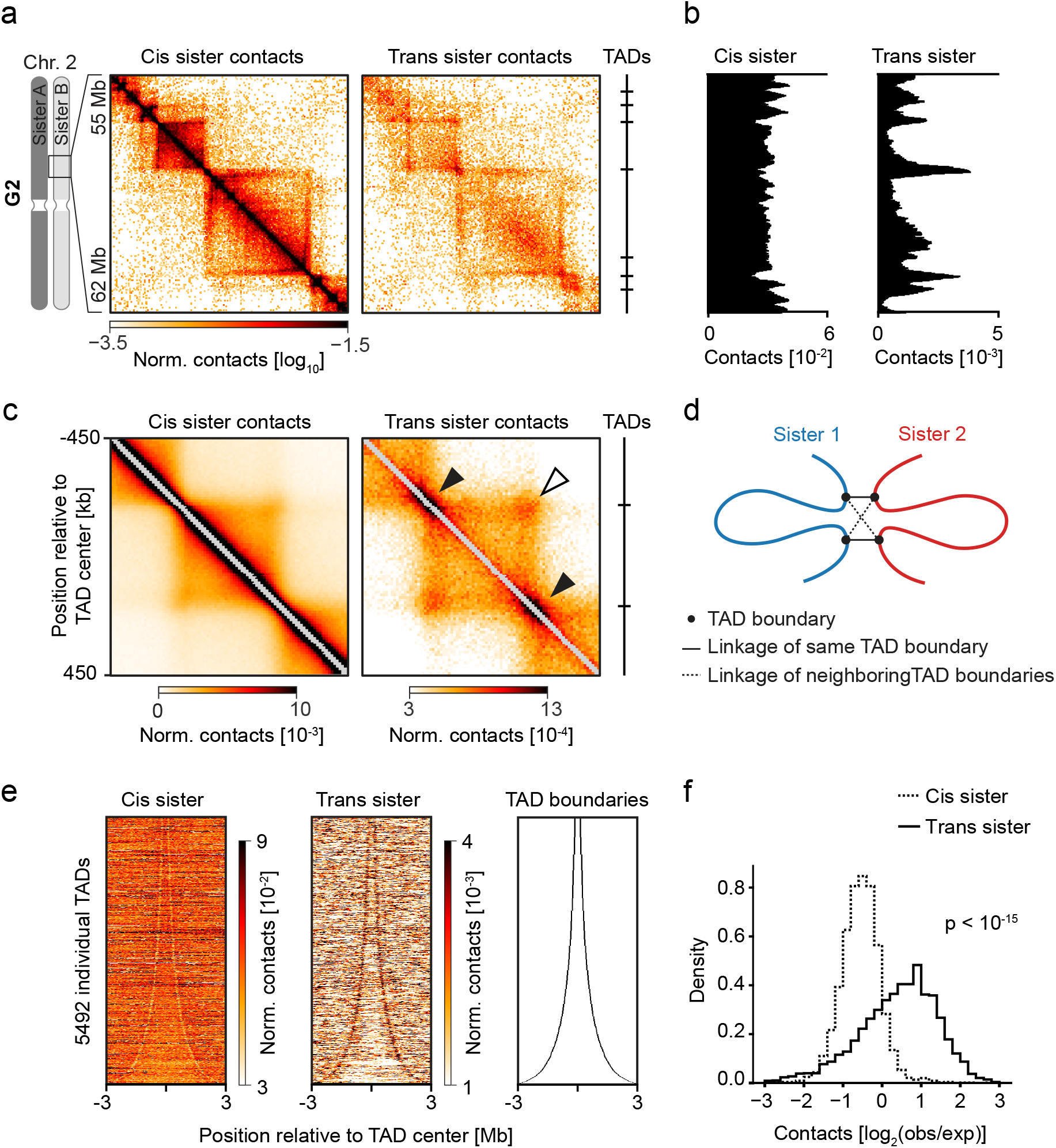
TAD topologies in replicated chromosomes. (a) Cis sister and trans sister contacts of 11 merged G2 samples at a representative region on chromosome 1 are displayed alongside the location of TAD boundaries (see Method section for details). Bin size of the matrix is 40kb. (b) Average trans sister and cis sister contact amount (“contact density”; see Methods for details) within a sliding window of 200 kb at the region shown in (a). (c) Average cis sister and trans sister contact environment around TAD centers of TADs between 300 and 500kb. The panel shows ICE-normalized contacts binned at 10 kb. Filled arrows indicate positions where the same TAD boundaries are connected across sister chromatids, whereas the hollow arrow indicates the connection of neighboring TAD boundaries across sister chromatids. (d) Model of sister-chromatid configuration around TAD boundaries. (e) Stack-up of average trans sister and cis sister contacts within sliding windows of 100kb along TADs sorted by size. The panel shows windows of 6 Mb around the center of the respective TADs.(f) Quantification of trans-contact enrichment at TAD boundaries. The average observed/expected values for cis sister and trans sister contacts within a 80 kb window surrounding all annotated TAD-boundaries (see Method section for details) are displayed as a histogram. P-value was calculated using a two-sided T-test.

## Molecular control of sister-chromatid topologies

The conformation of TADs in replicated chromosomes is organized by at least two functionally distinct types of cohesin complexes. During G2, about half of all chromatin-bound cohesin dynamically turns over^44,45^ to form cis-chromatid loops that shape TADs^6,16^, whereas the other half binds the stabilizing factor Sororin^27,46^ and persistently links sister chromatids. Trans sister contacts might concentrate at TAD boundaries because of motor-driven loop extrusion^20,21^ or via a mechanism involving cohesin independently of DNA loops. To investigate these possibilities, we aimed to selectively deplete the pool of cohesin that forms chromatin loops without disrupting sister-chromatid cohesion. The cohesin loading factor NIPBL is required to extrude and maintain cis-chromatid loops formed by cohesin^15,20^, but it is not expected to be required to maintain cohesion, given that this is mediated by persistently bound cohesin^44^. To test this hypothesis, we homozygously tagged NIPBL with auxin-inducible degrons (AID)^47^, synchronized cells to G2 and added auxin to induce NIPBL degradation (Extended Data Fig. 7a, b). Conventional Hi-C analysis showed a reduction of contact probability between ~200 kb and 3 Mb (Extended Data Fig. 7c), indicating suppression of cohesin-mediated loop formation. To assess whether cells maintain sister-chromatid cohesion under these conditions, we imaged live cells during mitosis. NIPBL-degraded cells efficiently congressed chromosomes (Extended Data Fig. 7d), confirming the presence of functional cohesion. Thus, NIPBL degradation during G2 selectively removes the pool of cohesin that forms loops and TADs while maintaining the pool of cohesin that mediates cohesion.

To determine how loop-forming cohesin affects sister-chromatid organization in G2 cells, we analyzed NIPBL-depleted cells by scsHi-C. TAD structures were much less pronounced and cis sister contacts were substantially reduced between ~200 kb and 3 Mb (Fig. 4a-c; Extended Data Fig. 7e, f, Table S3). Trans sister contacts were overall much more abundant compared to unperturbed control cells and particularly enriched along the diagonal throughout entire TADs (Fig. 4a-c, compare Fig. 3a-c), suggesting that sister chromatids generally interact more frequently in the absence of loop-forming cohesin. Analysis of individual genomic neighborhoods showed that there was less local enrichment of trans sister contacts around TAD boundaries (Fig. 4d) even though the overall number was greatly increased at TAD boundaries (Fig. 4e). Overall, these data suggest that the pool of cohesin that dynamically forms intra-chromatid loops is necessary to separate sister chromatids within TADs, resulting in locally enriched sister chromatid contacts at TAD boundaries.

**Figure 4.**
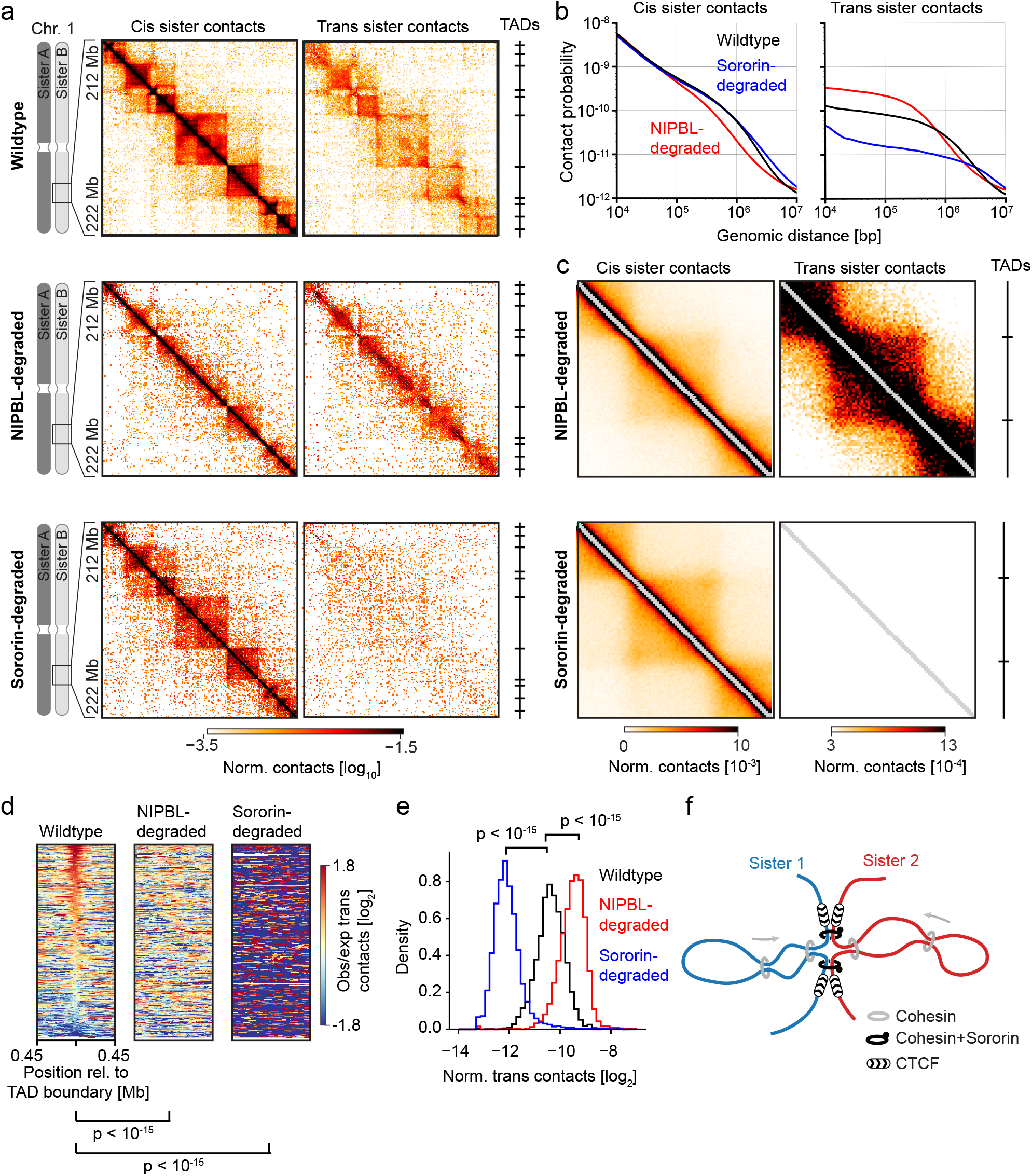
Organization of sister chromatids by distinct pools of cohesin complexes. (a) Cis sister and trans sister contacts of 11 merged G2 wildtype samples, 4 merged G2 NIPBL-degraded samples and 3 merged Sororin-degraded samples at a representative region on chromosome 1 are displayed alongside the location of TAD boundaries (see Method section for details). Bin size of the matrix is 40kb. (b) Average contact probability over different genomic distances for cis sister and trans sister contacts of the different G2 sample shown in (a). (c) Average cis sister and trans sister contact environment around TAD centers of TADs between 300 and 500kb in G2 synchronized cells with either degraded NIPBL or Sororin. The panel shows ICE-normalized contacts binned at 10 kb in a window of 900 kb. (d) Stack-up of average observed/expected values within sliding windows of 100kb around TAD boundaries of G2 wildtype samples, G2 NIPBL-degraded samples and Sororin-degraded samples. The panel shows windows of 900 kb. The rows are sorted based on the center enrichment of the G2 wildtype condition. P-values were calculated using a Mann-Whitney U test performed on the values in the center column of the respective stack-up matrix. (e) Quantification of trans sister contact density at TAD boundaries. The average ICE-corrected trans sister contact amount within a 400kb window surrounding all annotated TAD boundaries (see Methods section for details) is shown for G2 wildtype samples, G2 NIPBL-degraded samples and Sororin-degraded samples. P-values were calculated using a Mann-Whitney U test. (f) Model of sister chromatid organization by two distinct cohesin complexes.

We next addressed how the pool of cohesin mediating sister chromatid cohesion affects the conformation of replicated chromosomes. To this end, we suppressed establishment of cohesion while maintaining the pool of cohesin that forms loops by acutely degrading Sororin prior to DNA replication in cells in which endogenous Sororin was homozygously tagged with AID (Extended Data Fig. 7a). We then released cells to the subsequent G2 and performed scsHi-C (Extended Data Fig. 7b,c; Table S4). In Sororin-depleted cells, trans sister contacts were globally reduced and not enriched along the diagonal anymore (Fig. 4a, b, Extended Data Fig. 7e), indicating a complete loss of global sister chromatid alignment. Consistently, trans sister contacts were not enriched at TAD boundaries (Fig. 4d, e). However, the cis sister contact distribution was indistinguishable from that of unperturbed cells (Fig. 4a-c; Extended Data Fig. 7e). The Sororin-stabilized pool of cohesin is thus not required to form intra-chromatid loops or TADs in G2, but it is required to prevent the separation of sister chromatids and to maintain their global alignment during G2.

## Conclusions

Our high-resolution maps of replicated human chromosomes show that a pool of cohesin mediating linkage between replicated DNA molecules globally aligns sister chromatids during G2, while a pool of cohesin that dynamically forms loops locally separates sister chromatids within the confines of TAD boundaries (Fig. 4f). This organization has implications for the maintenance, expression, and mechanical transport of the genome.

The global alignment of sister chromatid arms by Sororin-stabilized cohesin favors interactions between homologous genome regions, as required for error-free homology-directed DNA damage repair ^13^. Local variations of sister chromatid pairing along chromosome arms might explain how the genomic context affects the efficiency of homology-directed DNA repair^48^. TAD boundaries are prone to DNA breakage, which can lead to chromosomal rearrangements^49,50^. Concentration of sister chromatid linkages at TAD boundaries might facilitate homology-directed repair of such breaks and thus contribute to the maintenance of chromosomal integrity. Tight sister chromatid pairing observed in TADs containing facultative heterochromatin might facilitate transcriptional co-repression and transfer of epigenetic information between distinct DNA molecules, as previously observed between paired homologous chromosomes in *D. melanogaster* ^51^, thereby contributing to the re-establishment of gene-regulatory domains after DNA replication. Conversely, separation of sister chromatids within TADs by loop-extruding cohesin might counteract trans-activation between promoters and enhancers on different DNA molecules, thereby improving the consistency of transcriptional output during cell cycle progression.

When cells enter mitosis, they resolve whole chromosome arms almost completely to enable sister subsequent chromatid segregation by the mitotic spindle. This involves sequential binding of two distinct condensin complexes and dissociation of cohesin from chromosome arms^9,52–54^, but how these activities are coordinated to promote sister chromatid resolution remains unknown. ScsHi-C provides a versatile tool to investigate this complex topological reorganization, as well as interactions between DNA molecules in other biological contexts, such as pairing and recombination of homologous chromosomes during meiosis.

## Acknowledgments

The authors acknowledge technical support by the IMBA/ IMP/GMI BioOptics and Molecular Biology Services facilities, and the Vienna BioCenter Metabolomics and Next Generation Sequencing facilities. We thank I. Patten, P. Batty, M.W.G. Schneider, M. Voichek for comments on the manuscript, and Life Science Editors for editing assistance. Research in the laboratory of D.W.G. is supported by the Austrian Academy of Sciences, an ERC Starting (Consolidator) Grant (nr. 281198), by the Wiener Wissenschafts-, Forschungs-und Technologiefonds (WWTF; project nr. LS14-009 and LS17-003), and by the Austrian Science Fund (FWF special research program “Chromosome Dynamics” SFB F34-06 and “RNAdeco” SFB F8011 and F8002, Doktoratskolleg “Chromosome Dynamics” DK W1238), Austrian Science Fund (stand alone projects P27947, P31691 to R.M), Austrian Research Promotion Agency FFG (West-Austrian BioNMR 858017 to R.M.). M.M. received a PhD fellowship from the Boehringer Ingelheim Fonds. The VBCF Metabolomics Facility is funded by the City of Vienna through the Vienna Business Agency. Research in the laboratory of J.-M.P. is supported by Boehringer Ingelheim, the Austrian Science Fund (FWF special research program SFB F34 “Chromosome Dynamics”), the Austrian Research Promotion Agency (Headquarter grant FFG-852936), the European Research Council (ERC) under the European Union’s Horizon 2020 research and innovation programme GA No 693949, and the Human Frontiers Science Programme (RGP0057/2018). Research in the laboratory of S.L.A. is supported by the European Research Council (ERC-StG-338252 and ERC-PoC-825710).

## Author contributions

Conceptualization, M.M., D.W.G.; Methodology, M.M., C.G., E.N., R.M., S.L.A.; Software, M.M., C.C.H.L.; Formal Analysis, M.M., C.C.H.L, A.G.; Investigation, M.M., Z.T., C.T.B.; Resources, W.T., G.J., J.M.P.; Writing, M.M., D.W.G., R.M.; Visualization, M.M.; Supervision D.W.G, R.M., J.M.P.; Funding Acquisition, D.W.G., J.M.P., R.M., S.L.A.

## Additional information

**Supplementary information** is available for this paper.

**Correspondence and requests for materials**should be addressed D.W.G.

---

**Extended Data Figure 1.**
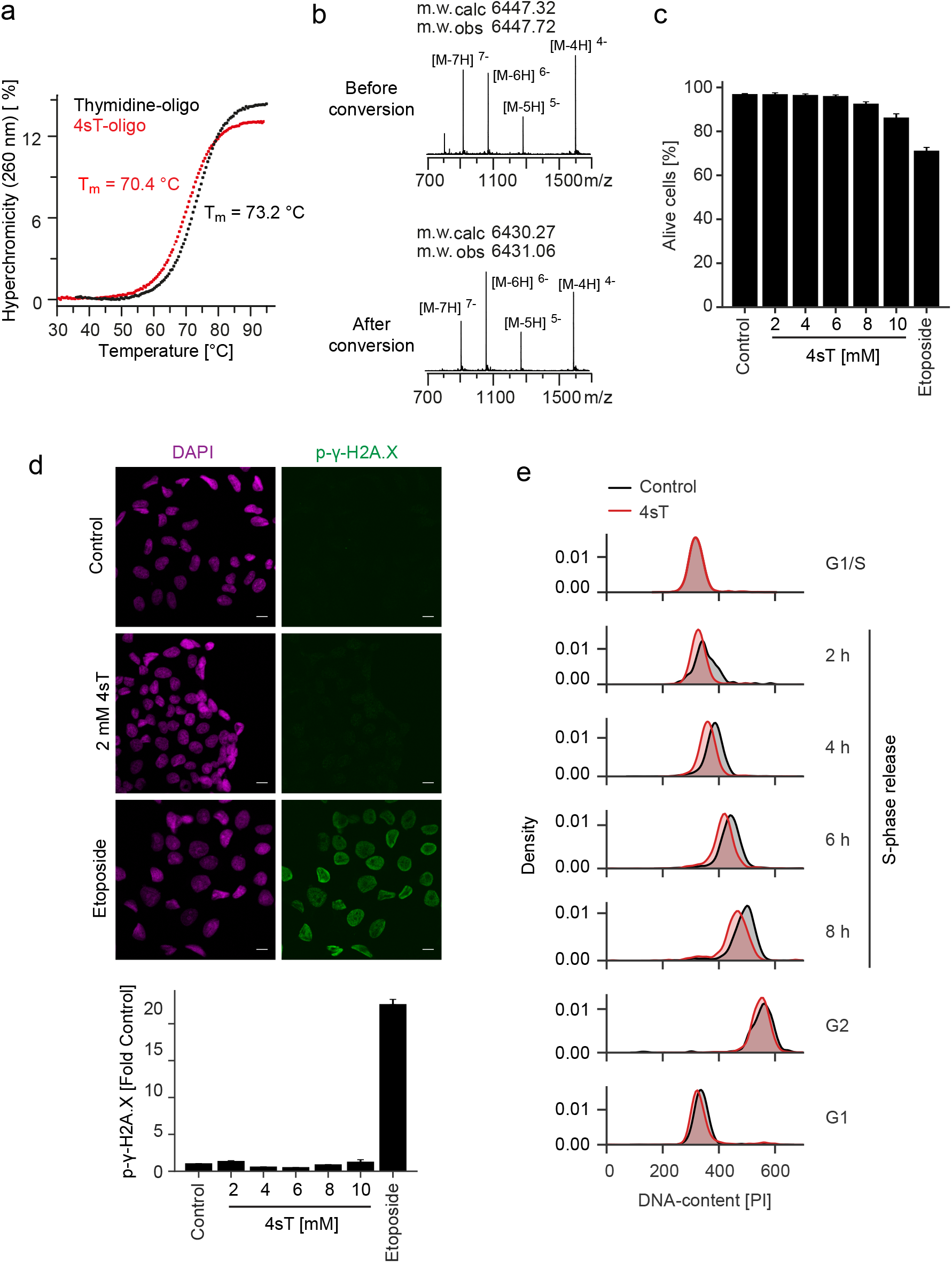
Characterization of 4sT. (a) Melting point analysis of a synthetic DNA hairpin containing 4sT or thymidine in the Watson–Crick base-paired stem, with sequence as depicted in Figure 1d (see Material and Methods for details). (b) Native mass spectrum of the oligonucleotide shown in Fig. 1d before and after OsO_4_/NH_4_Cl conversion measured in negative ion mode. (c) Percentage of live cells determined via Topro-3-Iodide staining of dead cells after 24 h incubation with the indicated compounds. Bars indicate mean and 95% confidence interval. (d) DNA damage assay performed after 24 h incubation with the indicated compounds. Quantification of mean fluorescence in cell nuclei stained by anti-p-γ-H2A.X antibody. Bars indicate mean and 95% confidence interval. Panel shows two pooled replicates. (e) Flow cytometry (FACS) analysis of cells progressing through S-phase in the presence or absence of 2 mM 4sT. DNA was stained using propidium iodide and kernel density estimation of signal in the PE-channel is shown. Cells were pre-synchronized to G1/S by thymidine and released into S-phase by removal of thymidine. The G2 sample was arrested by RO3306 and the G1 sample was arrested after progression through mitosis using thymidine.

**Extended Data Figure 2.**
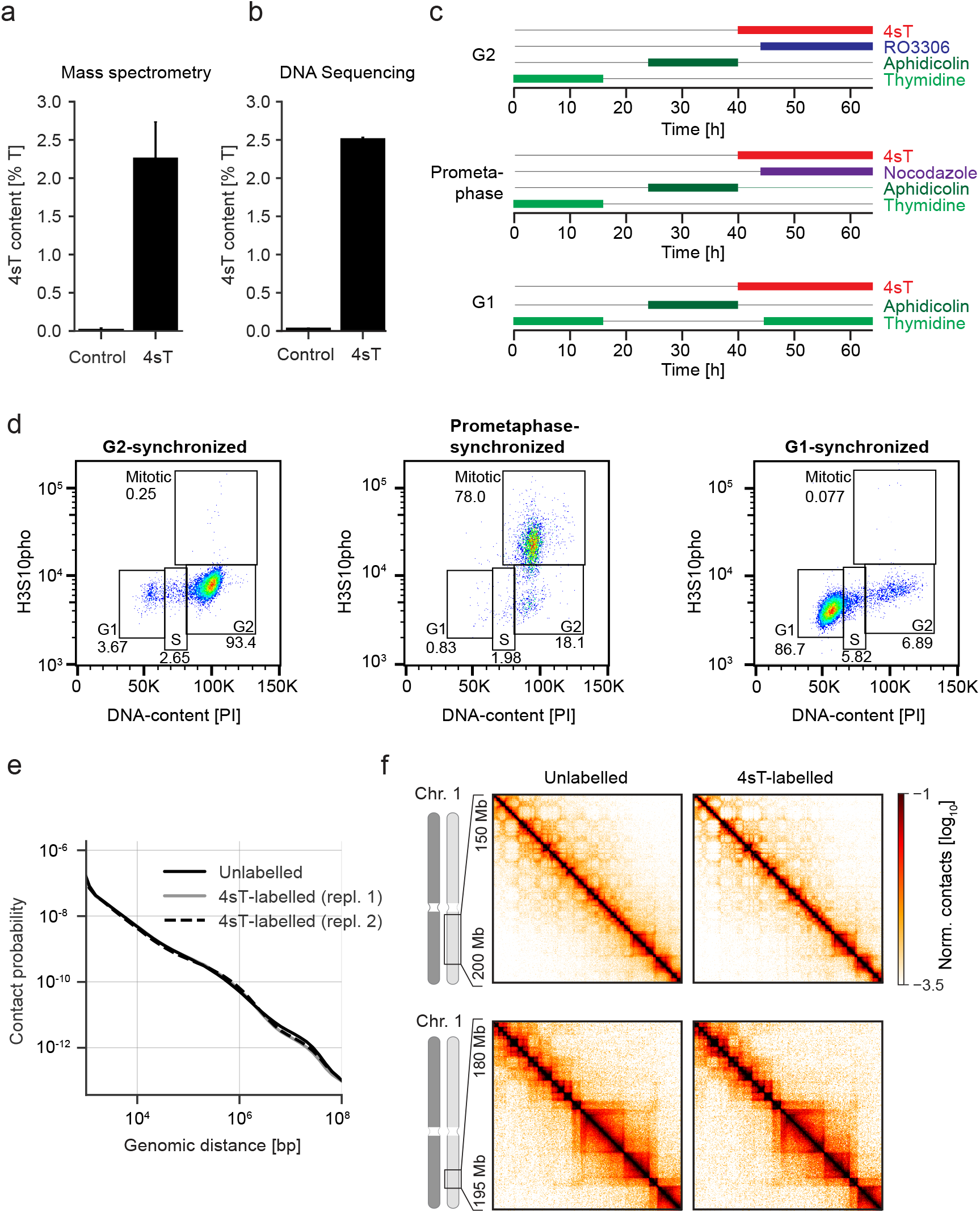
Preparation of cell cycle stage-specific scsHi-C samples. (a) Quantification of 4sT incorporation into genomic DNA of HeLa cells using mass spectrometry. Cells were grown in medium containing 4sT for 5 days and purified genomic DNA digested to single nucleotides. Indicated values reflect the percentage of 4sT in total measured thymidine. Bars indicate mean and 95% confidence interval of 3 independent experiments. (b) Quantification of 4sT incorporation using DNA sequencing. Cells were 4sT labelled as in (a) and purified genomic DNA chemically converted as in Fig. 1c. Indicated values are the sum of the A-to-G-mutation rate and the T-to-C mutation rate, normalized to the total amount of adenosine and thymidine measured respectively. Bars indicate mean and 95% confidence interval of 3 independent experiments. (c) Procedure to generate scsHi-C samples of cells synchronized to G2, prometaphase and the subsequent G1 phase. The different compounds were added to the cell culture medium as indicated by the colored bars. (d) Cell cycle analysis of WT HeLa cells synchronized to G2, prometaphase and G1 as indicated in (c). Anti-pH3S10 antibody was used to detect the mitotic state and propidium iodide to measure DNA content. Gates for different cell cycle stages are shown and the indicated numbers reflect percentage of cells that were measured. (e) Average contact probability over different genomic distances of HeLa cells synchronized to G2 that were either labelled with 4sT or unlabelled. (f) Hi-C interaction matrices at example regions of HeLa cells synchronized to G2 that were either labelled with 4sT or unlabelled.

**Extended Data Figure 3.**
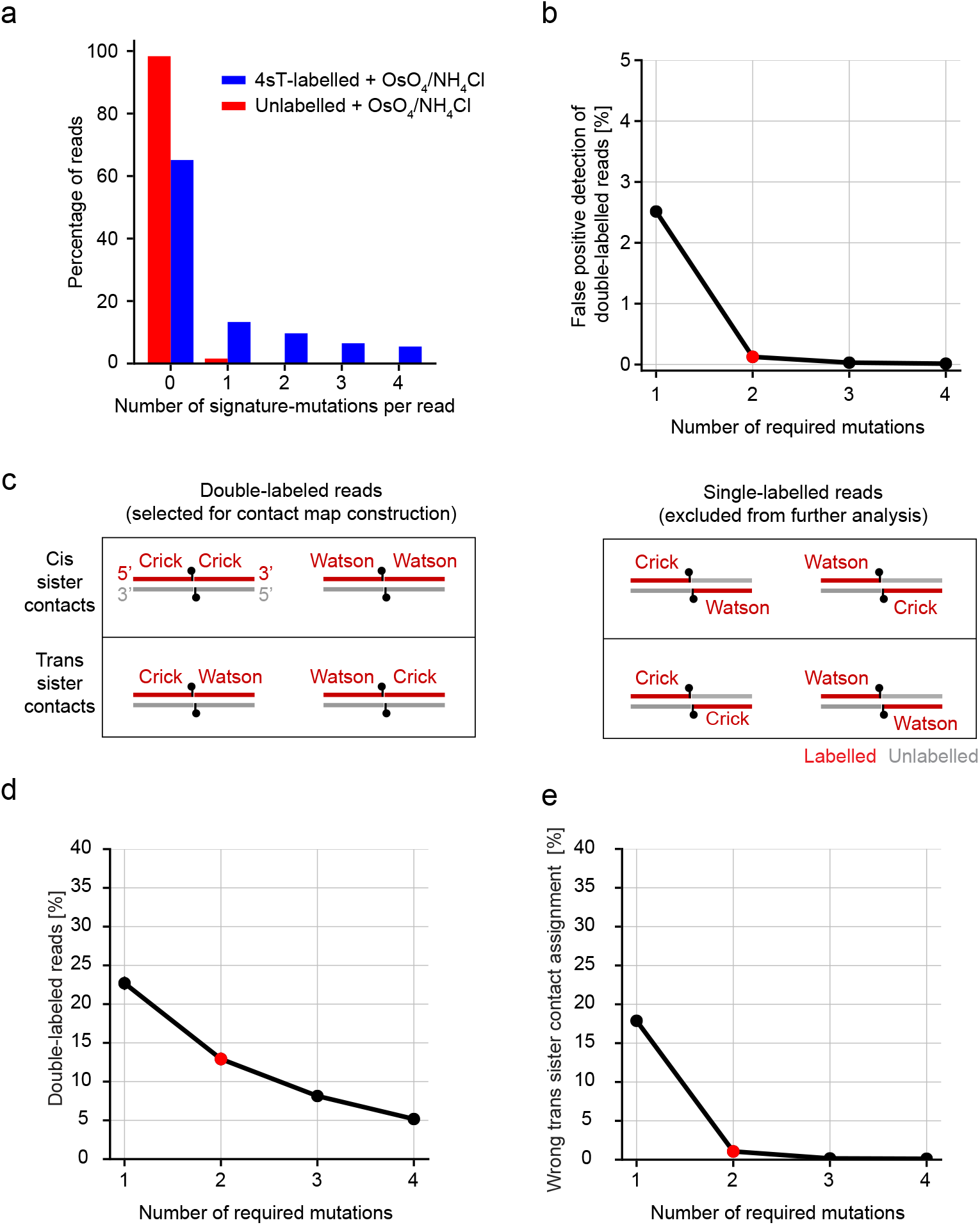
scsHi-C classification procedure. (a) Histograms of signature point mutations per read (AG or TC) of conventional sequencing libraries constructed from cells that were grown for 5 days in the presence (“4sT-labelled”) or absence (“unlabelled”) of 4sT and treated OsO_4_ as described in Fig. 1d. (b) False-positive rate of double-labelled read detection. Double-labelled reads were assigned based on different required signature point mutations on both read-halves. The sample shown was grown without 4sT, suggesting that every detected double-labelled read should be a false positive. (c) Depiction of all possible Hi-C ligation products of a sample where one strand of each sister chromatid has been labelled with a synthetic nucleotide. Only ligation products that carry two continuous halves that are labelled (“double-labelled”) can be used to discriminate cis sister contacts from trans sister contacts. This is because 4sT incorporation density is not high enough to allow detection of unlabelled reads based on the absence of signature mutations. If a given read exhibits signature mutations, however, it is possible to know with high confidence (see panel a and b) that it comes from the labelled strand. The ligation products that do not contain two halves with signature mutations are thus discarded during analysis. (d) Percentage of Hi-C contacts in HeLa wildtype cells synchronized to G2 in the presence of 4sT that can be used to assign sister chromatid identity (“double-labelled reads”) based on a classification scheme that requires more than the shown number of signature mutation thresholds. The number used in this paper (2) is highlighted in red. (e) Quantification of wrongly assigned trans sister contacts based on different signature mutation thresholds. To calculate the false-positive rate with which a cis sister contact is wrongly assigned as a trans sister contact, we adapted the scheme used to quantify the false-positive rate of trans-homolog contact assignment from Erceg et al^39^. Briefly, all contacts that exhibit a genomic separation smaller than 1kb are assumed to be Hi-C artifacts that arise from uncut continuous pieces of chromatin. Such contacts should be exclusively classified as cis sister contacts and thus all trans sister contacts in this range are assumed to be false positives. To then quantify the percentage of incorrect trans sister Hi-C contacts among all trans sister Hi-C contacts, the calculated false-positive rate was multiplied by the number of cis sister contacts exhibiting separation larger than 1kb and the resulting percentage of all trans sister contacts exhibiting separation larger than 1 kb plotted in this figure.

**Extended Data Figure 4.**
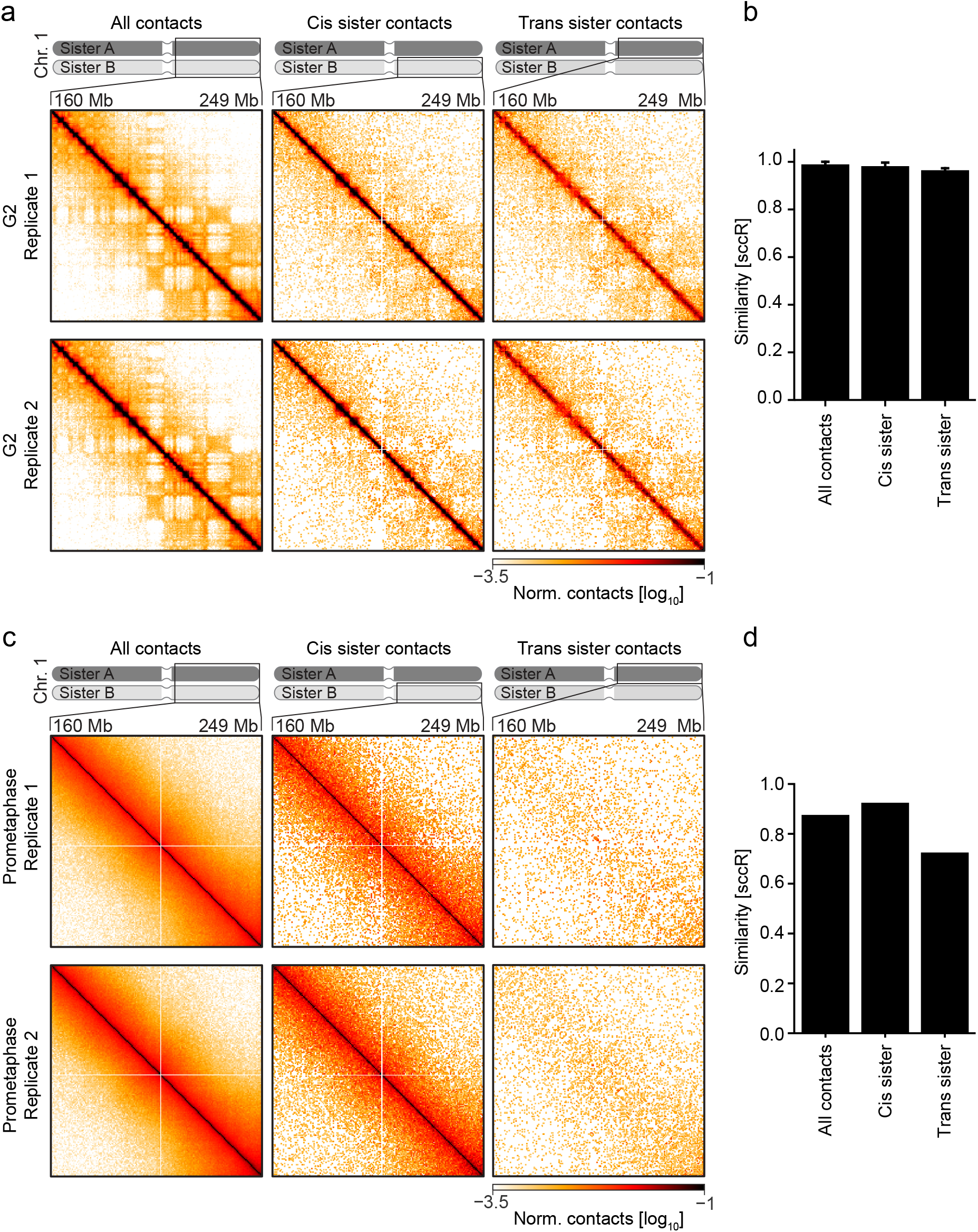
Reproducibility of scsHi-C. (a) Hi-C interaction matrices of the long arm of chromosome 1 of all contacts, cis sister, and trans sister contacts shown for two of the 11 G2 WT replicates. The all-contacts matrix was normalized to the total number of corrected contacts in the region of interest (ROI), whereas cis sister and trans sister contacts were normalized to the total amount of cis sister contacts and trans sister contacts in the ROI. Bin size of the matrix is 500 kb. (b) HiCrep^55^ analysis of all, cis sister and trans sister contacts of all 11 G2 replicates. Bars show the mean of all comparisons and the error shows the standard deviation. (c) Hi-C interaction matrix of the long arm of chromosome 1 of all, cis sister, and trans sister contacts of the two prometaphase replicates. Contacts were normalized as in (a). (d) HiCrep^55^ analysis of all, cis sister and trans sister contacts of two prometaphase replicates. Bars show the mean of all comparisons.

**Extended Data Figure 5.**
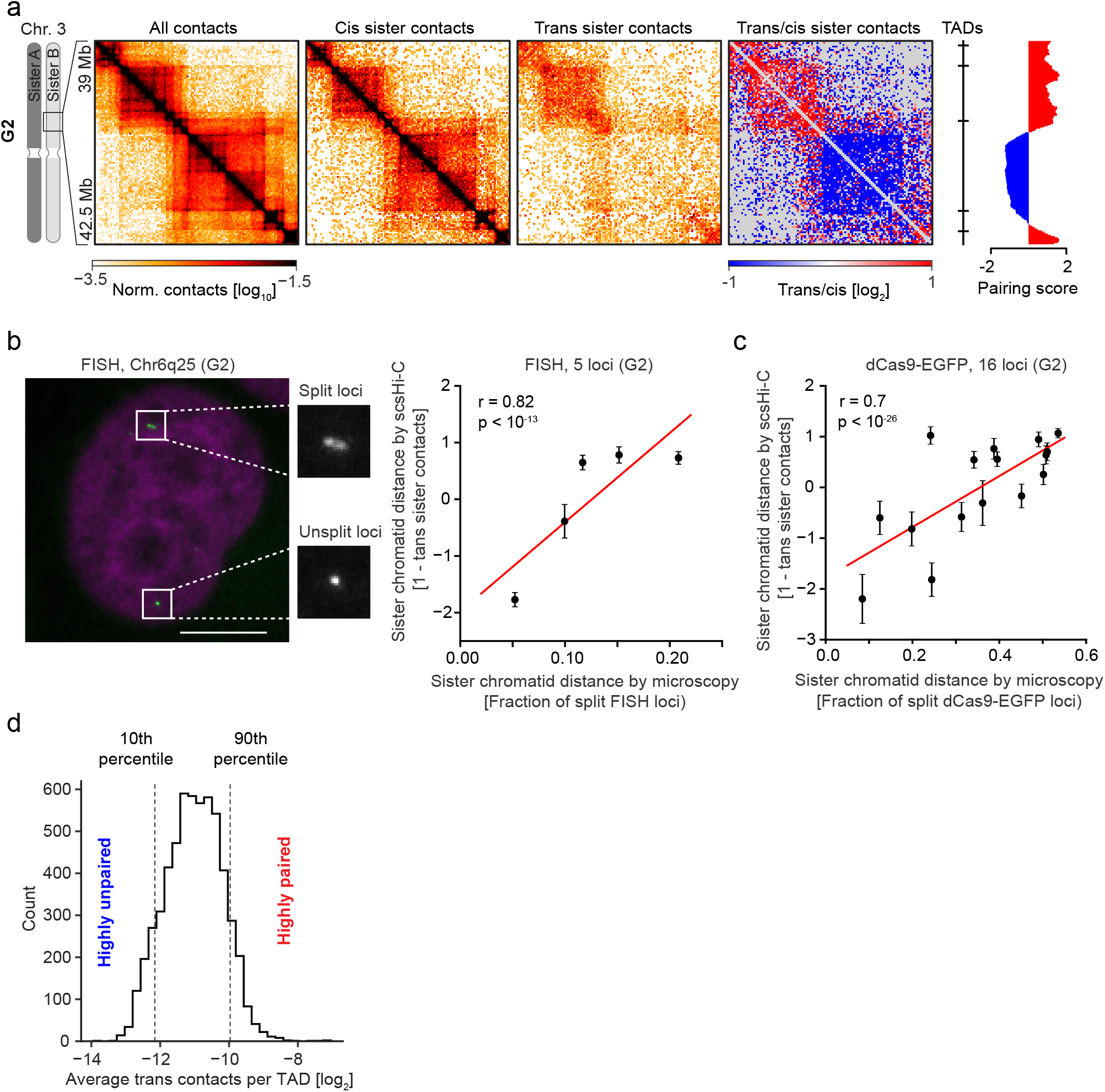
Sister chromatid conformation analysis by scsHi-C and microscopy. (a) All contacts, cis sister and trans sister contacts, as well as the ratio of trans sister observed/expected to cis sister observed/expected of 11 merged G2 samples at a representative region on chromosome 3 is displayed alongside the location of TAD boundaries and the trans sister pairing score (see Method section for details). Bin size is 30 kb. (b) Comparison of sister chromatid separation at 5 genomic loci measured by fluorescence in situ hybridization (FISH) and scsHi-C. Microscopy image shows examples for split and unsplit genomic sister loci, from G2-synchronized HeLa cell data reported in Stanyte et al^40^. scsHi-C quantification of sister locus distance was done by calculating (1 – average trans sister contacts) in a region spanning 600 kb around each FISH target site and standardizing the resulting value. Each dot indicates one target locus, measured in 11 independent HeLa WT G2 samples by scsHi-C. The error indicates the standard deviation of the Hi-C measurements. (c) Comparison of sister chromatid separation at 16 genomic loci measured by live cell microscopy and scsHi-C. Microscopy analysis was by live-cell imaging of 16 HeLa cell lines expressing dCas9-EGFP with different locus-specific gRNAs, using automated detection of merged or split sister loci in G2 cells, as reported in Stanyte et al^40^. scsHi-C quantification of sister locus distance was done by calculating (1 – average trans sister contacts) in a region spanning 600 kb around each gRNA target site and standardizing the resulting value. Each dot indicates one target locus, measured in 11 independent HeLa WT G2 samples by scsHi-C. The error indicates the standard deviation of the Hi-C measurements. (d) Histogram of average trans sister contact frequency for annotated TADs (see Method section for details). Vertical lines indicate the cut-offs for “highly paired” and “highly unpaired” TADs.

**Extended Data Figure 6.**
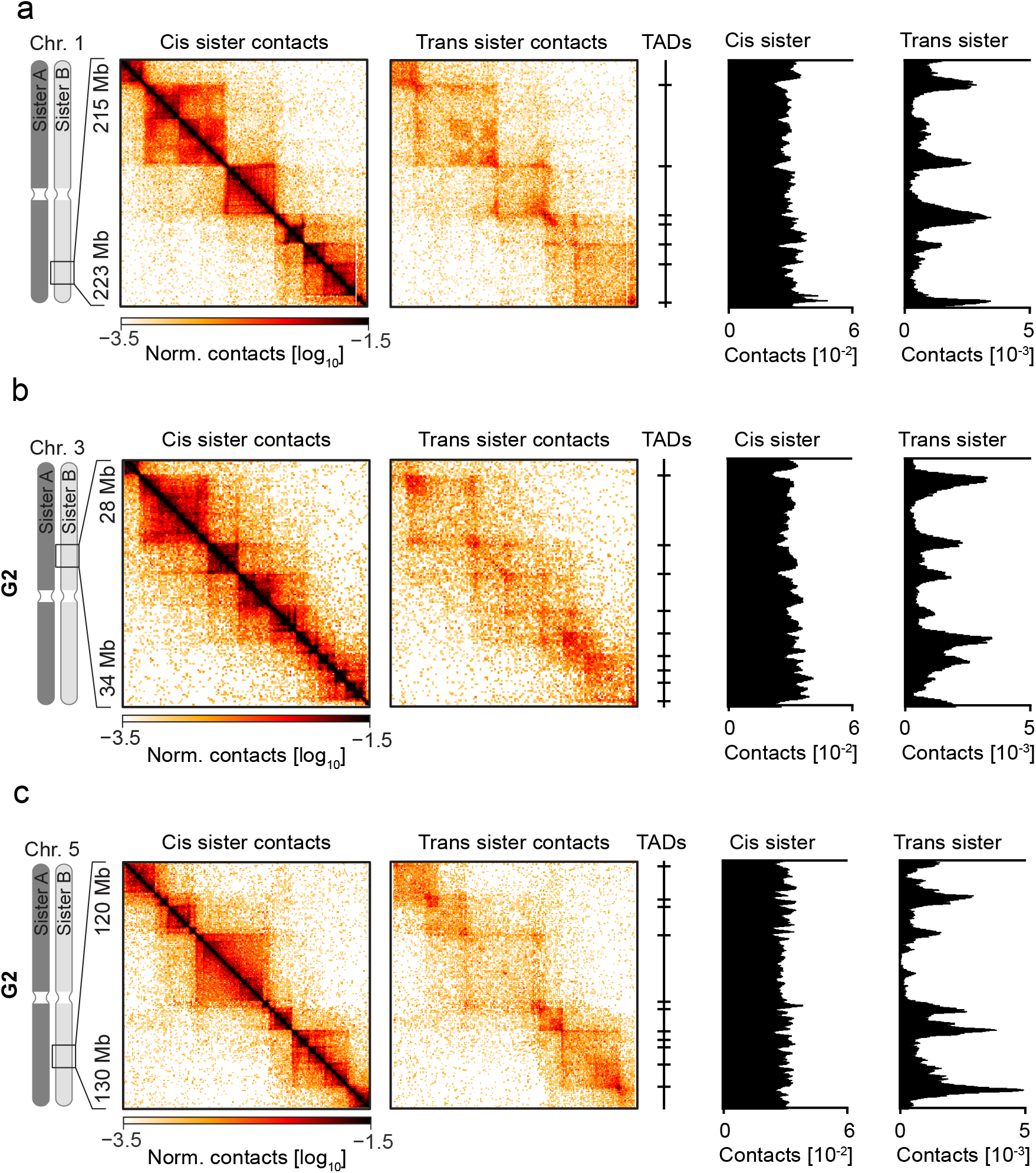
Sister chromatids are linked at TAD-boundaries. (a) Cis-sister- and trans-sister-contacts of 11 merged G2 samples at a representative region on chromosome 2 is displayed, alongside the location of TAD-boundaries (see Method section for details) and average trans-sister- and cis-sister-contact amount within a sliding window of 100 kb (see Method section for details). (b) Cis-sister- and trans-sister-contacts of 11 merged G2 samples at a representative region on chromosome 3 is displayed, alongside the location of TAD-boundaries (see Method section for details) and average trans-sister- and cis-sister-contact amount within a sliding window of 100 kb (see Method section for details). (c) Cis-sister- and trans-sister-contacts of 11 merged G2 samples at a representative region on chromosome 5 is displayed, alongside the location of TAD-boundaries (see Method section for details) and average trans-sister- and cis-sister-contact amount within a sliding window of 100 kb (see Method section for details).

**Extended Data Figure 7.**
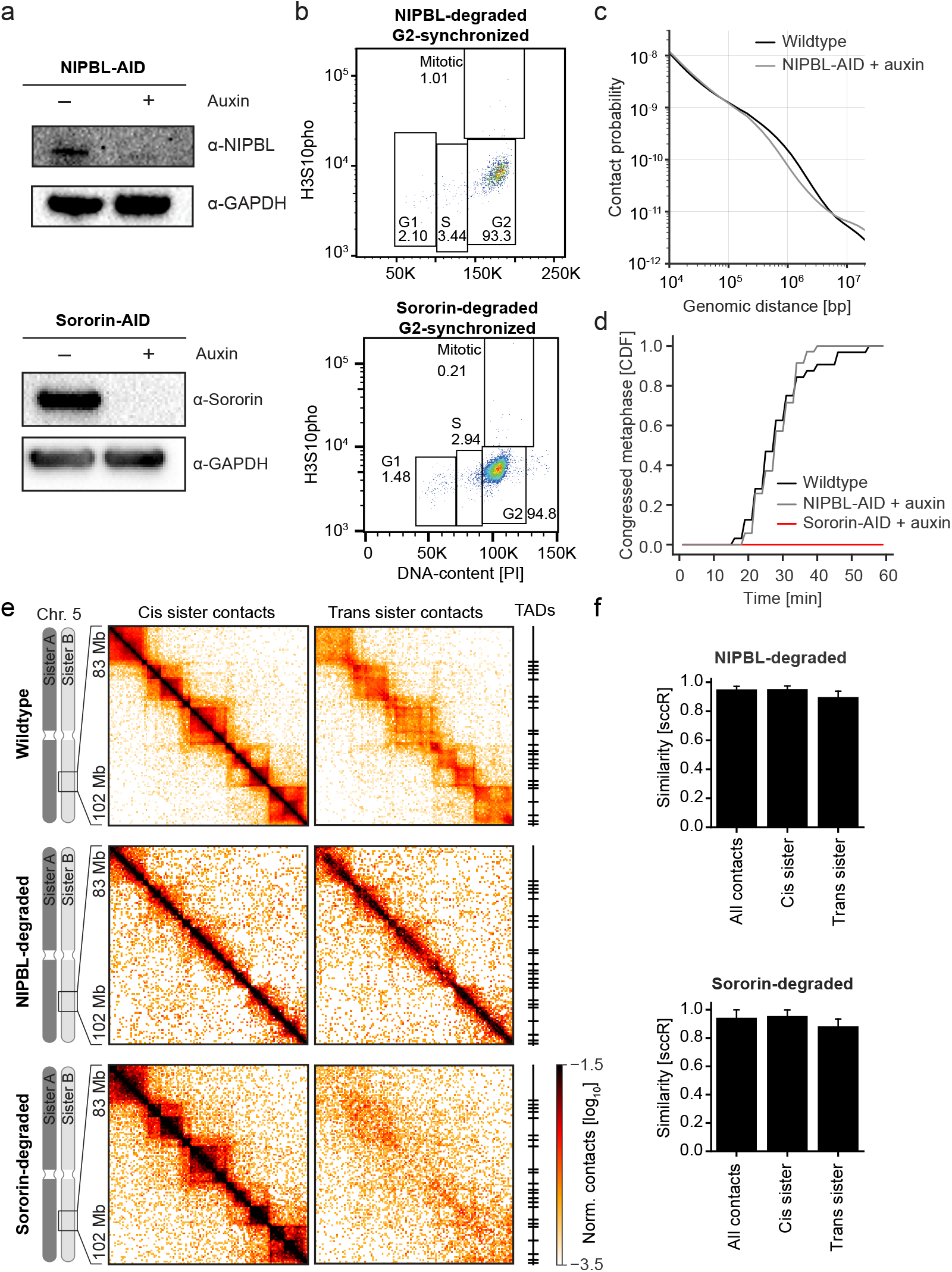
Characterization of HeLa Sororin-AID and HeLa NIPBL-AID cells. (a) Western blot for Sororin and GAPDH of HeLa Sororin-AID cells synchronized to G2 and either treated with auxin (+) or H_2_0 (-) as well as Western blot for NIPBL and GAPDH of HeLa NIPBL-AID cells synchronized to G2 and either treated with auxin (+) or H_2_O (-). (b) Cell cycle analysis of HeLa Sororin-AID and HeLa NIPBL-AID cells synchronized to G2 as indicated in Extended Data Fig. 2d, treated with auxin. Panel shows a FACS plot of cells stained for pH3S10 to mark mitotic cells and propidium iodide to measure DNA content. Gates for different cell cycle stages are shown and the indicated numbers reflect percentage of cells that were measured. (c) Contact probability of all contacts at different genomic distances of HeLa NIPBL-AID cells synchronized to G2 that were treated with auxin and HeLa WT cells synchronized to G2. (d) Metaphase congression analysis by time-lapse microscopy of WT HeLa cells, HeLa Sororin-AID cells and HeLa NIPBL-AID cells stained with SiR-DNA. HeLa Sororin-AID cells were treated with auxin before the final S-phase and HeLa NIPBL-AID cells were treated with auxin after the final S-phase. Panel shows the cumulative frequency of cells congressing their chromosomes in metaphase after entering mitosis in a RO3306 wash-out. Two pooled replicates are shown. (e) Cis sister and trans sister contacts of 11 merged G2 wildtype samples, 4 merged G2 NIPBL-degraded samples and 3 merged Sororin-degraded samples at a representative region on chromosome 5 are displayed alongside the location of TAD boundaries (see Method section for details). Bin size of matrix is 150kb. (f) HiCrep^55^ analysis of all, cis sister and trans sister contacts of all replicates of HeLa NIPBL-AID or Sororin-AID cells treated with auxin. Bars show the mean of all comparisons and the error shows the 95% confidence interval.

**Extended Data Figure 8.**
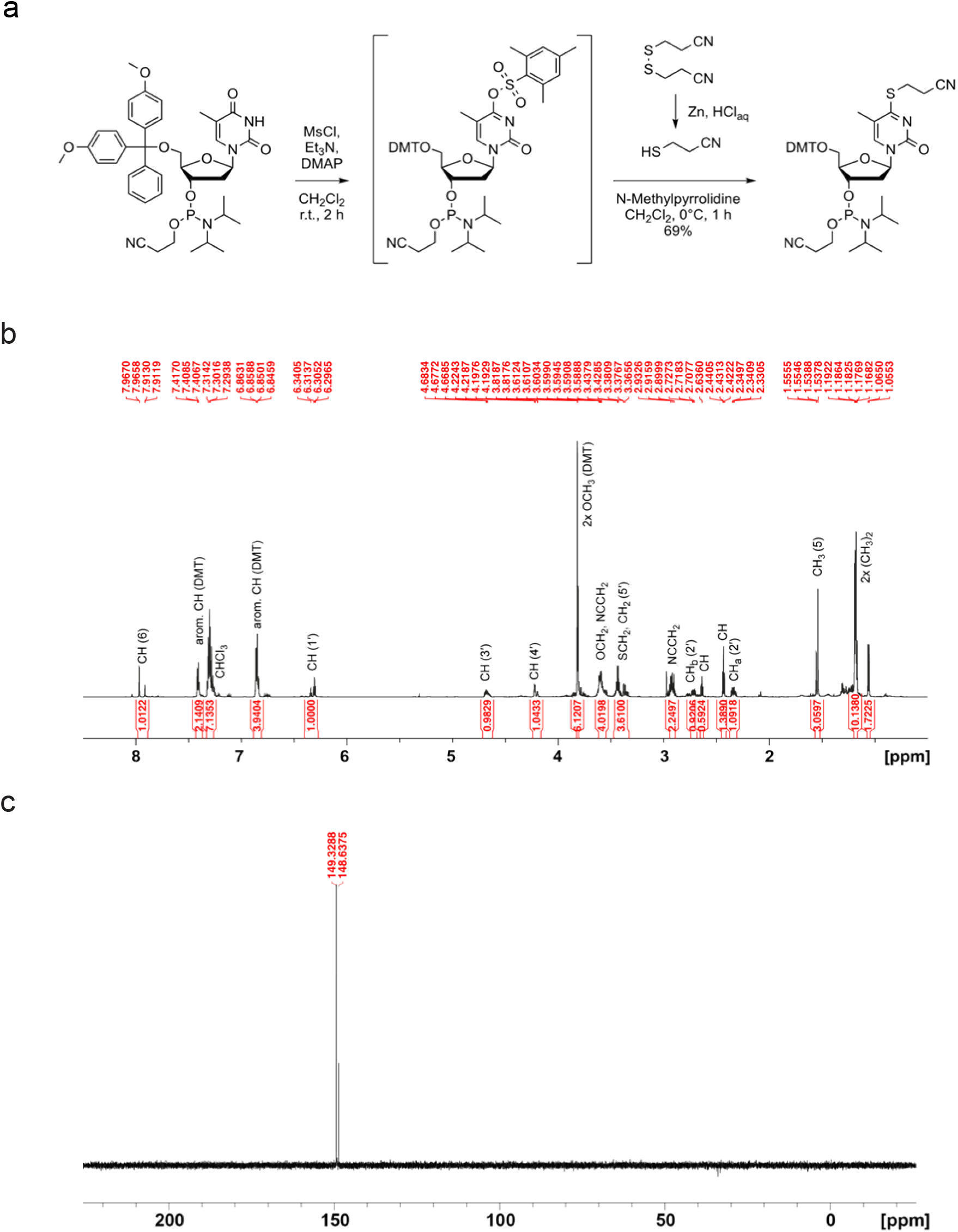
Synthesis of a 4sT-phosphoramidite building block. (a) Synthesis of 5′-*O*-(4,4′-Dimethoxytrityl)-*S*-(2-cyanoethyl)-4-thiothymidine 3′-*O*-[(2-cyanoethyl)-(*N*,*N*-diisopropyl)]-phosphoramidite. (b) ^1^H-NMR (700 MHz, CDCl_3_) of 4sT phosphoramidite (diastereomeric mixture). (c) ^31^P-NMR (282 MHz, CDCl_3_) of 4sT phosphoramidite (diastereomeric mixture).

## Materials and methods

### Cell culture

All cell lines used in this study have been regularly tested negatively for mycoplasm contamination. The parental HeLa cell line (‘Kyoto strain’) was obtained from S. Narumiya (Kyoto University, Japan) and validated by a Multiplex human Cell line Authentication test (MCA). Cells were cultured in WT medium (DMEM high-glucose [Sigma], buffered with HEPES [Applichem] and Sodium bicarbonate [Sigma], adjusted to pH 7.1-7.3 and supplemented with 10 % [v/v] FCS [Gibco], 1 % [v/v] Penicillin/Streptomycin [Gibco] and 1 % [v/v] GlutaMAX [Gibco]) in a humidified incubator at 37 °C and 5 % CO_2_. For culturing HeLa Sororin-AID and HeLa NIPBL-AID cells, medium was supplemented with 0.5 μg/ml Puromycin (Calbiochem). Cells were passaged every 48 h by dissociation using Trypsin/EDTA-Solution (Gibco).

### Generation of cell lines

All cell lines used in this study are listed in Table S6 and the plasmid used in their generation are listed in Table S7. The HeLa Kyoto N-terminally-tagged Sororin auxin-inducible degron (AID) cell line was created by CRISPR/Cas9-mediated genome editing as described previously ^16^. The gRNA sequences that were used for generating EGFP-AID-Sororin were CACCGCGCTCACCGGAGCGCTGAG, and CACCGACGTGAGGTCGAGCCGTTT together with the repair template ‘EGFP-AID-Sororin-HR’. The primers used for genotyping were CTGCGGGGGACAATACCAAT and CCGATCTCAGATTCCTGCCC. Subsequently, Tir1 expression was introduced by transducing a homozygous cell clone with lentiviruses using pRRL containing the constitutive promotor from spleen focus forming virus (SFFV) followed by *Oryza sativa* Tir1-3xMyc-T2A-Puro. Cells expressing Tir1 were selected by culturing in medium containing 2.5 μg/ml Puromycin (Sigma-Aldrich).

HeLa Kyoto N-terminally tagged AID-GFP-NIPBL cells were generated using CRISPR/Cas9-mediated genome editing based on a double nickase strategy ^56^. The template for homologous recombination introduced sequences coding for monomeric EGFP (L221K) and the *Arabidopsis thaliana* IAA17 71-114 (AID*) mini-degron ^57^. The gRNAs used for CRISPR/Cas9-mediated genome editing were CACCGCCCATTCATCCTGAATTTC and CACCGCCCCATTACTACTCTTGCG together with the repair template ‘AID-GFP-NIPBL-HR’. Single clones were obtained by sorting into 96-well plates on a BD FACS Aria III machine (BD Biosciences) and homozygous tagging was confirmed by PCR using the forward primer ATCGTGGGAACGTGCTTTGGA and reverse primer GCTCAGCCTCAATAGGTACCAACA. Subsequently, Tir1 expression was introduced as for HeLa Sororin-AID described above.

HeLa Kyoto RIEP H2B-mCherry cells were derived from HeLa Kyoto RIEP cells^58^ using a lentiviral delivery system^58^ to stably integrate a plasmid carrying H2B-mCherry (Lenti-H2B-mCherry). Cells were sorted into 96-wells to derive single clones using a BD Aria III instrument (BD Biosciences).

### FACS analysis of cell cycle stage

Cells were trypsinized, washed with phosphate buffered saline (PBS; made in-house) and fixed using 70 % EtOH (Sigma) for at least 1 h at 4 °C. Cells were spun down (1100 x g; 1 min) and permeabilized using 0.25 % Triton-X100 (Sigma) in PBS for 15 min on ice. Cells were spun down again and stained using 0.25 μg *α*-H3S10p (Merck Millipore 04-817) in 1 % Bovine serum albumin (BSA; Sigma) for 1 h at room temperature (RT). Cells were washed once with 1 % BSA and then stained using 1:300 *α*-mouse-AF488 (Molecular Probes A11001) in 1 % BSA for 30 min at RT in the dark. Cells were washed once with 1 % BSA and incubated with a solution containing 200 μg/ml RNase A (Qiagen) and 50 μg/ml Propodium Iodide (Sigma) in PBS for 30 min at RT in the dark. Samples were then measured on a FACSCanto instrument (BD Biosiences). Analysis was performed using FlowJo(v10) as follows: Gate for cells in FSC-A/SSC-A, for single cells in FSC-A/SSC-H, scatterplot of FITC and PI intensity.

### DNA damage assay

WT HeLa Kyoto cells were seeded into an 8-well Lab-Tek (Thermo Scientific) and grown for 16 h. Then, different concentrations of 4sT (2mM-10mM) and 50 μM etoposide (Sigma) were added and cells were incubated for 24 h. For immunofluorescence (IF), cells were washed two times with PBS and fixed using 4 % formaldehyde (Sigma) in PBS for 5 min. Formaldehyde was quenched using 20 mM TRIS-HCl (Sigma; adjusted to pH 7.5) in PBS for 3 min and washed with PBS. Cells were permeabilized using 0.5 % Triton-X100 (Sigma) in PBS for 10 min. Then, cells were blocked using 2 % BSA in PBS for 30 min at RT, followed by incubation with 1:500 *α*-phospho-*γ*-H2A.X (ABCAM ab2893) in 2 % BSA [PBS] for 1.5 h at RT. Then, cells were washed 3x for 5 min using PBS, followed by incubation with 1:1000 *α*-mouse-AF488 (Molecular Probes A11001) in 2 % BSA [PBS] for 30 min at RT in the dark. Then, cells were washed one time using PBS for 5 min, followed by staining using 1 μg/ml 4,6-diamidino-2-phenylindole (DAPI; Thermo Scientific) for 5 min. Then, cells were washed again for 5 min in PBS. Samples were imaged on a customized Zeiss LSM780 microscope using a 20x, 0.8 NA, Oil DIC Plan-Apochromat objective (Zeiss). Images were analyzed using CellCognitionExplorer ^59^ for segmentation and intensity extraction and Python scripts to visualize the data.

### Viability assay

Cells carrying a stable H2B-mCherry integration were seeded into a 96 well imaging plate (Greiner) in imaging medium (custom; DMEM High-glucose [Gibco] without Riboflavin and Phenolred containing 10% [w/w] FCS [Gibco], 1% [w/w] P/S [Gibco] and 1 % [w/w] Glutamax [Gibco]) supplemented with 1 μM TO-PRO^®^-3 (Thermo Fisher Scientific). After 16 h, compounds to be tested were added and imaging was started on a Molecular Devices ImageXpressMicro XL screening microscope with a reflection-based laser auto focus and a 10x, 0.75 NA, S Fluor dry objective (Nikon). Cells were maintained for 24 h in a microscopic stage incubator at 37 °C in a humidified atmosphere at 5 % CO_2_ and images in the mCherry and TO-PRO^®^-3 channel were recorded every 2 h. Images were analyzed using CellCognition ^60^ for segmentation and intensity extraction and Python scripts to visualize the data.

### Western Blot

Cell suspension (1 million cells/ml) was mixed with 6x SDS loading buffer and 10 mM DTT (Roche) and incubated at 95 °C for 10 min. Samples were separated on a NuPage 4-12% Bis-Tris Gel (Invitrogen) and transferred onto a Hybond P 0.45 polyvinylidene difluoride (PVDF) membrane (GE life sciences) using wet blotting. Sororin was probed using a custom antibody kindly provided by Jan-Michael Peters (1:500). GAPDH was probed using a polyclonal antibody (Abcam ab9485) and NIPBL was probed using a monoclonal antibody (Absea 010702F01). Horseradish peroxidase-conjugated secondary antibodies were used, and blots were visualized using ECL Plus Western Blotting Substrate (Thermo Fisher Scientific).

### Metaphase congression assay

Cells were synchronized to G2 as explained in the cell synchronization section (Extended Data Fig. 2c). Then, 1 h before RO3306 wash-out, cells were supplemented with 250 ng/ml SIR-DNA (Spirochrome). Then, cells were washed 2x with imaging medium containing 250 ng/ml SIR-DNA, followed by imaging every 3 min for 120 min on a customized Zeiss LSM780 microscope at 37 °C and 5 % CO_2_ using a 20x, 0.8 NA, Oil DIC Plan-Apochromat objective (Zeiss). Congression time was measured by visual inspection using Fiji ^61^ for all cells that entered mitosis in the indicated time frame. Results were visualized using Python.

### Cell synchronization for scsHi-C

Cells were seeded into 25 cm^2^ flasks and grown for 3 h, then supplemented with 2 mM Thymidine (Sigma). Cells were released 16 h later by washing 2 times with prewarmed WT medium. 8 h later, cells were supplemented with 3 μg/ml aphidicolin (Sigma) and 2 mM 4sT. Cells were released 16 h later by washing 2 times with PBS and addition of medium containing 2 mM 4sT. For Sororin-AID experiments, 500 μM Indole-3-acetic acid (Sigma) was added 1 h prior to S-phase release. For S-phase release experiments, samples were taken at the indicated time-points. For synchronization to G2 and prometaphase, after 4 h release, 9 μg RO-3306 (Sigma) or 200 ng/ml nocodazole (Sigma) were added respectively. For NIPBL-AID experiments, 500 μM Indole-3-acetic acid (Sigma) was added 8 h after released. Samples were processed 16 h later. Cells were harvested by washing with PBS, followed by trypsinisation and resuspension in WT medium. Cells were then spun down, washed again with PBS, followed by fixation for 4 min in 1 % formaldehyde (Sigma). Cell pellets were stored at −20 °C or processed immediately.

### Hi-C sample preparation

Fixed cells were permeabilized using ice-cold Hi-C lysis buffer (10 mM TRIS-HCl pH 8 [Sigma], 10 mM NaCl [Sigma], 0.2 % Nonidet P-40 substitute [Sigma], 1x Complete EDTA-free Protease inhibitor [Roche]) for 30 min at 4 °C. Then, cells were spun down (2500 x g for 5 min), supernatant was discarded, and digestion mix was added (375 U DpnII [NEB] in 1x DpnII buffer [NEB]) and cells incubated for 16 h at 37 °C under rotation. Then, cells were spun down, supernatant was discarded and fill-in mix was added (38 μM Biotin-14-dATP [Thermo Fisher Scientific], 38 μM dCTP, dGTP and dCTP [Thermo Fisher Scientific], 50 U Klenow Polymerase [NEB], 1x NEB 2 buffer) and cells incubated for 1 h at 37 °C under rotation. Then, cells were spun down again, and ligation mix was added (1x T4 DNA ligase buffer [Thermo Fisher Scientific], 0.1 % Triton X-100 [Sigma], 100 μg/ml BSA [Sigma], 50 U T4 DNA ligase [Thermo Fisher Scientific]) and incubated at RT for 4 h. Then, cells were spun down, resuspended in 200 μl PBS and gDNA was purified using the DNeasy Blood and Tissue kit (Quiagen). DNA was transferred to a Covaris microTUBE (Covaris) and sheared on a Covaris S2 instrument (Duty cycle 10 %, Intensity 5.0, Cycles/burst 200) for 25 s. Double size selection was performed by employing AMPure XP beads (Beckman Coulter) first at 0.8-fold sample volume according to the standard protocol, followed by transfer of the supernatant and bead application at 0.12-fold sample volume. The resulting DNA was bound to Dynabeads MyOne Streptavidin C1 beads (Thermo Fisher Scientific) in Biotin binding buffer (5 mM Tris-HCl pH 7.5 [Sigma], 0.5 mM EDTA [AppliChem], 1 M NaCl [Merck]) for 1 h at RT. Beads were then washed 2x in Tween wash buffer (5 mM Tris-HCl [Sigma], 0.5 mM EDTA [AppliChem], 1 M NaCl [Merck], 0.05 % Tween-20 [Sigma]) and 1x in H_2_O. Beads were resuspended in H_2_O and library preparation was performed using the NEBNext Ultra II DNA library prep kit for Illumina [NEB] according to the standard protocol. After this, beads were washed 4x using Tween wash buffer and DNA was eluted using 95 % formamide (Sigma), 10 mM EDTA (AppliChem) at 65 °C for 2 min. DNA was then precipitated using 80 % EtOH (Sigma), washed with 75 % EtOH and resuspended in H_2_O. Then, 4sT was converted to methyl-cytosine using OsO_4_ / NH_4_Cl (see below), followed by qPCR according the NEBUltra Ultra II DNA library prep kit for Illumina [NEB]. The finished libraries were purified using AMPure XP beads (Beckman Coulter) at 0.9x sample volume following the standard protocol.

### Sequencing

Sequencing of all samples was performed either on an Illumina NovaSeq instrument using patterned SP flowcells using read-mode PE250 or on an Illumina MiSeq instrument using a Nanoflowcell using read-mode PE300 (v2).

### Quantification of 4sT incorporation into gDNA

Deoxyribonucleosides were quantified by injecting 1 μl of the acidified digest on a RSLC ultimate 3000 (Thermo Fisher Scientific) directly coupled to a TSQ Vantage mass spectrometer (Thermo Fisher Scientific) via electrospray ionization. A Kinetex C18 column was used (100 Å, 150 x 2.1 mm), employing a flow rate of 100 μl/min. An 8-minute-long linear gradient was used from 0% A (1 % acetonitrile, 0.1 % formic acid in water) to 60% B (0.1 % formic acid in acetonitrile). Liquid chromatography-tandem mass spectrometry (LC-MS/MS) was performed by employing the selected reaction monitoring (SRM) mode of the instrument. Thymidine and 4-thio-thymidine were quantified by analyzing the in-source fragments of the respective nucleotides at an elevated declustering potential. For thymidine the transition 127.1 m/z → 54.1 m/z (CE 23 V) and for 4-thio-thymidine the transition 143.1 m/z → 126.1 m/z (CE 25 V) were used. A calibration curve of synthetic standard nucleosides was used to quantify the relative percentage of 4-thio-thymidine in total thymidine in the biological samples. Each sample was measured in duplicate.

### Conversion analysis of 4sT on synthetic oligos

A 4sT-containing oligonucleotide was synthesized as described below. The molecular weight of the oligonucleotide with and without OsO_4_ / NH_4_Cl treatment (see below) was analyzed on a Finnigan LCQ Advantage MAX ion trap instrument connected to an Amersham Ettan micro LC system in the negative-ion mode with a potential of −4 kV applied to the spray needle. LC: Sample (200 pmol RNA dissolved in 30 μl of 20 mM EDTA solution; average injection volume: 30 μl), column (Waters XTerra^®^ MS, C18 2.5 m; 2.1 × 50 mm) at 21 °C; flow rate: 30 μl min^−1^; eluent A: Et_3_N (8.6 mM), 1,1,1,3,3,3-hexafluoroisopropanol (100 mM) in H_2_O (pH 8.0); eluent B: MeOH; gradient: 0–100% B in A within 30 min; UV detection at 254 nm.

### Conventional sequencing library preparation to estimate 4sT mutation rates

Cells were harvested by trypsinization, spun down at 1100 x g for 1 min and the supernatant was discarded. Then, cells were spun down, resuspended in 200 μl PBS and gDNA was purified using the DNeasy Blood and Tissue kit (Quiagen). DNA was transferred to Covaris microTUBE (Covaris) and sheared on a Covaris S2 instrument (Duty cycle 10 %, Intensity 5.0, Cycles/burst 200) for 25 s. Double size selection was performed by employing AMPure XP beads (Beckman Coulter) first at 0.8-fold sample volume according to the standard protocol, followed by transfer of the supernatant and bead application at 0.12-fold sample volume. DNA library preparation was performed with the resulting DNA using the NEBNext Ultra II DNA library prep kit for Illumina [NEB] according to the standard protocol. The unamplified libraries were then treated using OsO_4_ (see below) and amplified according the NEBNext Ultra II DNA library prep kit for Illumina [NEB]. The finished libraries were purified using AMPure XP beads (Beckman Coulter) at 0.9x sample volume following the standard protocol.

### OsO_4_ / NH_4_Cl-mediated conversion of 4sT

For synthetic oligos, lyophilized DNA (1 nmol) was dissolved in water (10 μl) and denatured for 2 min at 90 °C. Then, the solution was heated to 60 °C and NH_4_Cl buffer (2 μl, 2 M, pH 8.88) and OsO_4_ solution (10 μl, 1 mM) were added to yield final concentrations of 0.45 mM OsO_4_ and 180 mM NH_4_Cl in a total volume of 22 μl. The reaction mixture was incubated for three hours at 60 °C. The DNA was precipitated by adding 90 μl of precipitation solution (made of water (650 μl), aqueous NaOAc solution (150 μl; 1 M, pH 5.2), and glycogen (10 μl; 20 mg/ml)) and 250 μl of cold ethanol. The mixture was kept at –20 °C for 30 minutes, followed by centrifugation (13,000 rpm, 4 °C, 30 min). The supernatant was discarded, and the precipitated DNA analyzed by anion exchange HPLC and mass spectrometry (see above). Genomic DNA was incubated with 0.45 mM OsO_4_ (Sigma) and 200 mM NH_4_Cl (Sigma) adjusted with NH_3_ (Honeywell Fluka) to pH 8.88 first for 5 min at 95 °C followed by 60 °C for 3 h on a T100 Thermal Cycler (Bio-Rad) with the heated lid set to 105 °C. DNA was then precipitated using 80 % EtOH, washed with 75 % EtOH and resuspended in H_2_O.

### Melting curve analysis of hairpin oligo

The absorbance versus temperature profiles were recorded at 260 nm on a Varian Cary 100 spectrophotometer equipped with a multiple cell holder and a Peltier temperature-control device. The 4sT containing DNA hairpins and their reference oligonucleotides were measured at a concentration of 2 μM in melting buffer (150 mM NaCl, 10 mM Na_2_HPO_4_ at pH 7.0). Three cycles of cooling and heating between 30 °C to 95 °C and a rate of 0.7 °C min^−1^ were recorded. Sample preparation: An aliquot of oligonucleotide stock solution was lyophilized, dissolved in 1 ml of melting buffer to give the desired final concentration. The solution was transferred into a quartz cuvette and degassed. A layer of silicon oil was placed on the surface of the solution to minimize evaporation during the measurements.

### Statistical analysis and sample number

All Hi-C datasets that are presented in this paper are merges of at least two independent replicates. For a detailed listing of all datasets, see Table S1-S4. All statistical tests were performed using scipy ^62^.

### Data Reporting

No statistical methods were used to predetermine sample size. The experiments were not randomized. The investigators were not blinded to allocation during experiments.

### Synthesis of 4-thiothymidine (4sT)-containing oligodeoxynucleotides

Synthesis of a 4sT-phosphoramidite building block; general information. Chemical reagents and solvents were purchased from commercial suppliers and used without further purification. Thymidine phosphoramidite was purchased from ChemGenes. Organic solvents for reactions were dried overnight over freshly activated molecular sieves (3 Å). All reactions were carried out under argon atmosphere. Analytical thin-layer chromatography (TLC) was carried out on Marchery-Nagel (Polygram SIL G/UV254, 0.2 mm silica gel) plates. Flash column chromatography was carried out on silica gel 60 (70-230 mesh). 1H and 31P NMR spectra were recorded on Bruker 400 and 700 MHz spectrometers. The chemical shifts are referenced to the residual proton signal of the deuterated solvents: CDCl3 (7.26 ppm), d6-DMSO (2.50 ppm) for 1H NMR spectra; 31P-shifts are relative to external 85% phosphoric acid. 1H assignments were based on COSY experiments. Mass spectrometric analysis of low molecular weight compounds was performed on a Thermo Scientific Q Exactive Orbitrap mass spectrometer in the positive ion mode. Procedure: Thymidine phosphoramidite (229 mg, 0.307 mmol) was dissolved in dry dichloromethane (3 ml). Then, triethylamine (37 mg, 51 μl, 0.366 mmol), 4-dimethylaminopyridine (1 mg) and 2-mesitylensulfonyl chloride (56 mg, 0.256 mmol) were added. The solution was stirred at room temperature for 2 hours. In the meantime, 3,3′-dithiobis(propionitrile) (200 mg, 1.16 mmol) was suspended in aqueous 2 M HCl solution, followed by slow addition of zinc powder (220 mg, 3.36 mmol). The mixture was stirred at room temperature for one hour, extracted three times with dichloromethane, dried over Na_2_SO_4_ and evaporated. The 3-mercaptopropionitrile was obtained as slightly yellow oil. Then, *N*-methylpyrrolidine (261 mg, 319 μl, 3.07 mmol) and the freshly prepared 3-mercaptopropionitrile (133 mg, 1.54 mmol) were mixed in dry dichloromethane (1 ml) and added to the reaction mixture containing the activated nucleoside. Stirring was continued at 0 °C (ice bath) for one hour. Finally, the solution was diluted with dichloromethane, washed with saturated NaHCO_3_, dried over Na_2_SO_4_ and evaporated (Extended Data Fig. 8a). The crude product was purified by column chromatography on silica gel (ethyl acetate/cyclohexane, 15:100 – 75:25). Yield: 172 mg (69%) white foam. TLC (ethyl acetate/cyclohexane, 1:1): R_f_ = 0.1. HR-ESI-MS (m/z): [M+Na]+ calculated: [836.3217]; found: [836.3110]. 1H-NMR (400 MHz, CDCl_3_): δ 1.15 – 1.18 (m, 12 H, 2x (H_3_C)_2_CHN); 1.43 (s, 3H, H_3_C(5)); 2.22 – 2.28 (1H, HaC(2′)); 2.33 – 2.35 (1H, HC(N)); 2.53 – 2.56 (1H, HC(N)); 2.60 – 2.66 (1H, HbC(2′)); 2.78 – 2.88 (2H, H_2_CCN); 3.26 – 3.34 (m, 4H, H_2_CS, H_2_C(5′)); 3.44 – 3.54 (m, 4H, H_2_CCN, H_2_CO); 3.73 (s, 6H, 2x H_3_CO(DMT)); 4.13 (m, 1H, HC(4′)); 4.58 (m, 1H, HC(3′)); 6.21 (m, 1H, HC(1′)); 6.73 – 6.78 (m, 4H, HC(DMT)); 7.27 – 7.41 (m, 9H, HC(DMT)); 7.87 (s, 1H, HC(6)) ppm. ^31^P-NMR (282 MHz, CDCl_3_): δ 148.64; 149.33 ppm (Extended Data Fig. 8b-c).

### Solid-phase synthesis of 4sT containing DNA

CCGGAAGGTATGAACC(4sT)TCCG was synthesized by automated solid-phase synthesis (ABI 392 Nucleic Acids Synthesizer) using standard DNA nucleoside phosphoramidites (ChemGenes), the 4-thiothymidine phosphoramidite (as described above), and polystyrene support (GE Healthcare, Primer Support 80s, 80 μmol per g; PS 200). The following set-up was applied: detritylation (80 s) with dichloroacetic acid/1,2-dichloroethane (4/96); coupling (2.0 min) with phosphoramidites/acetonitrile (0.1 M, 130 μl) and 5-(benzylthio)-1*H*-tetrazole/acetonitrile (0.3 M, 360 μl); capping (0.4 min, three cycles) with Cap A: 4-(dimethylamino)pyridine in acetonitrile (0.5 M) and Cap B: Ac_2_O/sym-collidine/acetonitrile (2/3/5); oxidation (1.0 min) with I_2_ (20 mM) in THF/pyridine/H_2_O (35/10/5). Acetonitrile (DNA synthesis grade) was purchased from Anteris Systems GmbH. Acetonitrile, acetonitrile solutions of amidites, and acetonitrile solution of 5-(benzylthio)-1*H*-tetrazole were dried over activated molecular sieves (3 Å) overnight.

### Deprotection of 4sT containing DNA

After DNA strand assembly, the beads were treated with 1,8-diazabicyclo[5.4.0]undec-7-en (DBU) in anhydrous acetonitrile (5 ml, 1 M) for three hours. Subsequently, the beads were incubated with *tert.*-butyl amine/ethanol/water (1/1/2, v/v/v) and dithiothreitol (50 mM) for five hours at 55 °C. Then, the supernatant was removed and the beads were washed three times with 1 ml ethanol/water (1/1). The combined phases were evaporated to dryness. The crude DNA was dissolved in water (1 ml).

### Analysis and purification of 4sT containing DNA

After the deprotection, the crude DNA was analyzed by anion-exchange chromatography on a Dionex DNAPac PA-100 column (4 mm × 250 mm) at 80 °C. Flow rate: 1 ml min^−1^, eluant A: 25 mM Tris·HCl (pH 8.0), 6 M urea; eluant B: 25 mM Tris·HCl (pH 8.0), 0.5 M NaClO_4_, 6 M urea; gradient: 0 – 60% B in A within 50 min, UV detection at 260 nm. The DNA was purified on a semipreparative Dionex DNAPac PA-100 column (9 mm × 250 mm) at 80 °C with flow rate 2 ml min-1, using the same eluents A and B as for analytical analysis, but with flat gradients that were optimized according to the length of the oligonucleotide. DNA containing fractions were loaded on a C18 SepPak Plus cartridge (Waters/Millipore), washed with 0.1 − 0.15 M (Et_3_NH)^+^HCO ^−^, H_2_O and eluted with H_2_O/CH_3_CN (1/1). DNA containing fractions were evaporated to dryness and then dissolved in 1 ml water (stock solutions for storage at –20 °C). The quality of purified DNA was again analyzed by anion-exchange chromatography. The molecular weight of the DNA was analyzed by LC-ESI MS. Yields were determined by UV photometrical analysis of oligonucleotide solutions.

## Sequencing data analysis

### Published datasets used

All published datasets used in the sequencing data analysis are listed in Table S5.

### Data and code availability

All datasets used in this study will be uploaded to GEO. The ipython notebooks used to perform all the sequencing data analysis of data generated within this work are available at https://github.com/gerlichlab/scsHiCanalysis, alongside with a detailed description of each script and the figures they produce. The environment used to perform this analysis is provided as a docker container (https://hub.docker.com/repository/docker/gerlichlab/scsHiC) under the tag “release-1.0”. All the versions of the software packages used are noted within the dockerfile.

### Calling HeLa single-nucleotide polymorphisms

In order to discriminate between 4sT-introduced mutations and single-nucleotide polymorphisms (SNPs), HeLa Kyoto SNPs were called on DNA-seq data from WT HeLa cells and a new consensus genome based on hg19 was constructed using bcftools (https://github.com/samtools/bcftools).

### Mutation rate analysis

First, sequencing data was aligned to the hg19 genome containing HeLa SNPs using bowtie2. Then, relative point mutation rates for all possible point mutations were calculated using a custom Python script as follows: Absolute point mutation rates were counted and normalized to the total covered amount of the source base (e.g. for T-to-C normalization was done to T in the reference genome). These values were then normalized to an unlabelled control sample that had undergone OsO_4_ / NH_4_Cl treatment. The incorporated amount of 4sT was determined by sequencing analysis, based on calculating the ratio of the sum of T-to-C and the A-to-G absolute mutation rates to the sum of all Ts and all As, respectively. The fraction of read pairs being labelled in samples derived from DNA-seq libraries was calculated as follows: Only high-quality read-pairs (alignment score > 20, Phred-score > 20, longer than 240 bp) were counted. Only point mutations that had a Phred-score higher than 20 were counted. The number of T-to-C and A-to-G mutations were counted on both read-pairs of a paired-end read. The read halves were then assigned to be labelled if they contained 2 or more signature point mutations. A read-pair was classified as labelled if any of the two halves were labelled. Rates of double labelled reads for Hi-C samples were calculated similarly, but a read-pair was only classified as labelled if both halves were labelled. To calculate a histogram of signature mutations per read, the number of T-to-C and A-to-G mutations of high-quality reads (see above) was counted and plotted for control samples (not treated with 4sT) and samples treated with 4sT for 5 days.

### Correlation analysis of loci splitting frequency and FISH with scsHi-C

The average number of trans sister contacts (balanced as described in “Hi-C data preprocessing”) was extracted at target sites of the 16 gRNAs from Stanyte et al. ^40^ within a 600 kb window and 1 − *average*(*trans* − *sister contacts*) was calculated and converted to a Z-score by subtracting the mean and dividing by the standard deviation. The resulting value correlated with the average frequency of split loci 1.2 h before G2 phase ^40^ for 11 WT G2 replicates. A similar analysis was performed for the 5 loci for which both gRNA splitting data and FISH data was available.

### Hi-C data preprocessing

Hi-C samples were preprocessed using a custom nextflow pipeline (https://github.com/gerlichlab/scshic_pipeline). Briefly, bcl2 files were first demultiplexed using bcl2tofastq. Then, fastq files were aligned to hg19 with HeLa SNPs using bwa, aligning read pairs independently. Then, pairsam files were constructed using pairtools (https://github.com/mirnylab/pairtools), followed by sorting and deduplicating. Then, reads were split into cis sister and trans sister contacts based on the presence of signature mutations using pairtools select: A read was assigned to the Watson strand if it contained two or more A-to-G mutations and no T-to-C mutations. Similarly, if a read contained two or more T-to-C mutations, but no A-to-G mutations it was assigned to the Crick strand. Then, contacts were classified as cis sister contacts if (after correcting for the opposite read-strandedness of Illumina sequencing of the two mates) both mates mapped to the same strand. Conversely, contacts were classified as trans sister contacts if the two mates mapped to opposing strands. Then, cooler (https://github.com/mirnylab/cooler; ^63^) was used to construct.cool files, and binned at multiple resolutions. After completion of the nextflow pipeline, cis sister and trans sister contacts were merged, and the resulting file balanced using cooltools ^64^. Balancing was done as described in ^64^, excluding the 0^th^-diagonal to avoid Hi-C artefacts. Bins that had marginal read-count with a median absolute deviation (MAD) > 5 based on the genome-wide distribution were excluded from balancing and further analysis. Then, the resulting weights were transferred to the individual cooler files containing the cis sister and trans sister contacts. Hi-C matrices containing all contacts (not stratified into cis sister and trans sister contacts) were balanced similarly.

### Hi-C genome scaling plots

Scaling plots were calculated separately for cis sister and trans sister contacts using pairlib (https://github.com/mirnylab/pairlib). Briefly, contacts were binned into geometrically spaced bins from 10 kb to 100 Mb with a total of 64 bins. Then, the number of contacts in each bin was divided by the number of covered base pairs. When multiple samples were compared on the same plot, they were down sampled to contain an equal number of combined cis sister- and trans sister contacts that are separated further than 1 kb using the NGS package (https://github.com/gerlichlab/NGS).

### Observed-over-expected transformation of Hi-C matrices

The expected number of Hi-C contacts *e* at a given genomic separation *k* in Hi-C bin units was calculated using the cooltools (https://github.com/mirnylab/cooltools) package:

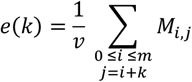

with *e*(*k*) being the expected number of Hi-C contacts separated by *k* Hi-C bins, *v* being the number of valid bin-interactions with separation *k* (interactions between bins that were assigned valid balancing weights during the ICE-procedure) and *M* being the ICE-corrected Hi-C interaction matrix containing *m* bins. Note that the expected number of contacts is obtained from the upper-triangular part of the Hi-C matrix only since the matrix is symmetric. The observed-over-expected Hi-C matrix *OE* was then obtained as follows:

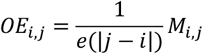

### Hi-C aggregate maps at TAD-centers

Aggregate maps of Hi-C submatrices around TAD-centers of genomic neighborhoods were calculated within a custom ipython notebook using the cooltools package (https://github.com/mirnylab/cooltools). First, 900 kb-sized submatrices centered around TAD-centers were extracted and the pixel-wise average of the ICE-corrected contacts over all the windows was calculated. In order to avoid Hi-C artefacts, the main diagonal as well as the neighboring diagonals were blanked out in the plot.

### Extraction of sample regions

Sample regions of ICE-corrected Hi-C-matrices were extracted using the cooler Python API. For calculations of ratio maps, the observed/expected values were calculated for the respective ROI using the cooltools (https://github.com/mirnylab/cooltools) package as described above and then trans/cis ratios were calculated. Before plotting, a pseudocount of 0.01 was added to avoid removal of 0-bins from the image during log-transformation.

### TAD-calling

TAD-calling was done using OnTAD^41^ on a G2 WT Hi-C matrix construct from all generated contacts merged over all replicates. The bin size for TAD-calling was 50kb and the only parameter that was changed from the standard set was the maximum TAD-size, which was restricted to 6 Mb. The TADs used for all analysis in this paper can be found here https://github.com/gerlichlab/scsHiCanalysis/blob/master/data/TADs_final.bedpe.

### Pairing-score and contact-density calculation

We defined the contact density as the average contact frequency within a sliding window of half-length *w* in Hi-C bin units:

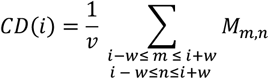

with *CD*(*i*) denoting the contact density at bin *i*, *M* the Hi-C matrix (either ICE-corrected or observed-over-expected transformed), *v* being the number of valid Hi-C pixels within the window of summation (interactions between bins that were assigned valid balancing weights during the ICE-procedure; Note that the main diagonal does not contain valid pixels) within the sliding window. We then defined the pairing-score to be the contact density subtracted by the genome-wide average and converted to a Z-score by dividing by the genome-wide standard deviation:

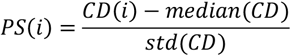

with *PS*(*i*) referring to the pairing score at genomic bin *i*, *median*(*CD*) referring to the genome-wide median of *CD* and *std*(*CD*) referring to the genome wide standard deviation of *CD*.

### Stack-up analysis of line profiles

Stack-ups of line profiles along a set of regions was calculated as follows:

1. The contact density within a sliding diamond of half-length *w* was calculated along each region within the set of regions as described above (either for ICE-corrected matrices or observed-expected-transformed matrices), resulting in a vector of size *n* for each region.
2. Then, these vectors were stacked into an *m* × *n* matrix with *m* denoting the number of regions and *n* denoting the length of the line profile along each region as described in 1.
3. Finally, the rows of the matrix were sorted based either on the size of TADs within the regions of interest for analyses in Fig.2f and Fig.3e or based on the average line profile signal within the center bins – bins with index in the interval [*median*(0, *n*)⌋ − 5, ⌊*median*(0, *n*)⌋ + 5] - for analysis in Fig.4d.

For display of observed-over-expected transformed values, a pseudocount of 0.01 was added before log-transformation. Moreover, line profiles that only contained invalid Hi-C bins were removed from the stack-up.

### LOLA-analysis of highly paired and highly unpaired TADs

LOLA ^42^ enrichment analysis was done for TADs with high trans sister intensity and low trans sister intensity as follows: The average contact density (see above for details) for every annotated TAD (see above for details) was calculated for a window with size *w* that corresponded to the size of the respective TAD centered on the TAD-center 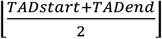 for a Hi-C contact matrix binned at 10kb and containing ICE-corrected trans sister contacts. The 90th and 10th percentile of trans sister contact density within these TADs was calculated and the TADs that showed a trans sister contact density larger than the 90th percentile were denoted “highly paired”, whereas TADs that had a trans sister contact density smaller than the 10th percentile were denoted “highly unpaired”. LOLA was then run using the LOLA Extended Dataset ^42^ for the highly paired and highly unpaired TAD regions using all TADs as the region universe. Only chromatin datasets that were from HeLa cells are shown.

### HiCRep analysis

HiCRep^55^ was run using the python wrapper *hicreppy* (https://github.com/cmdoret/hicreppy) for all conditions tested using a Hi-C matrix with bin size 100kb, a smoothing parameter *v* = 10, a maximum distance of *maxdist* = 10^6^*bp* without subsampling.

### False-positive rate estimation of double labelled reads and trans sister contacts

To estimate the false-positive rate of double labelled Hi-C contact, a Hi-C sample from cells that were grown in the absence of 4sT was analyzed. The reasoning was that all reads that were classified as double labelled in this condition would be false positives. A Hi-C contact was annotated as double labelled, if both half-reads exhibited more than a threshold amount of signature mutations (A to G or T to C). Then, the false-positive rate

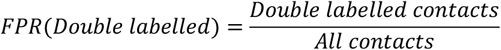

was calculated for different thresholds of signature mutations. To estimate the false-positive rate of trans sister contact assignment, an approach developed in^39^ was adapted. We assumed that all contacts of a G2 WT Hi-C sample that exhibited a genomic separation below 1kb were Hi-C artefacts, namely uncut DNA. Such contacts should be exclusively classified as cis sister contacts since a successful digestion and re-ligation is needed to generate a trans sister contact. The false-positive rate of trans sister contacts was therefore defined as

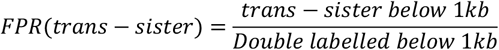

To estimate the number of wrongly assigned trans sister Hi-C contacts we assumed that the false-positive rate of trans sister assignment is independent of genomic separation and that the amount of wrongly assigned cis sister contacts is negligible. We further assumed that contacts with genomic separation larger than 1kb constitute true Hi-C contacts. We therefore defined the fraction of wrong trans sister Hi-C contacts *WTC* as

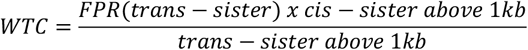

and calculated this value for different signature mutation thresholds.

### H3K27me3 enrichment analysis

In order to calculate the enrichment of H3K27me3 Chip-seq signal at highly paired and highly unpaired TADs, ChIP-seq data from ^65^ was downloaded and the average enrichment of H3K27me3 with respect to the control dataset calculated for both region sets.

---

**Table S1.** Read statistics of the G2 wild-type samples generated in this study.

|  | Total | Mapped | Unique | Unique (HQ) | Cis | Trans heterolog | Cis 1kb+ | Cis 10kb+ | Cis sister | Trans sister |
| --- | --- | --- | --- | --- | --- | --- | --- | --- | --- | --- |
| Replicate 1 | 3.08E+08 | 2.32E+08 | 1.00E+08 | 9.54E+07 | 9.35E+07 | 6.90E+06 | 4.26E+07 | 2.92E+07 | 1.26E+07 | 6.56E+05 |
| Replicate 2 | 3.55E+08 | 2.64E+08 | 1.06E+08 | 1.01E+08 | 9.91E+07 | 7.21E+06 | 4.43E+07 | 2.95E+07 | 1.29E+07 | 6.28E+05 |
| Replicate 3 | 2.24E+08 | 1.74E+08 | 1.28E+08 | 1.22E+08 | 1.20E+08 | 8.51E+06 | 4.86E+07 | 3.20E+07 | 1.62E+07 | 6.84E+05 |
| Replicate 4 | 3.57E+08 | 2.71E+08 | 1.46E+08 | 1.39E+08 | 1.34E+08 | 1.25E+07 | 6.28E+07 | 4.24E+07 | 1.72E+07 | 9.97E+05 |
| Replicate 5 | 3.37E+08 | 2.61E+08 | 1.40E+08 | 1.33E+08 | 1.28E+08 | 1.17E+07 | 5.80E+07 | 3.92E+07 | 1.76E+07 | 9.56E+05 |
| Replicate 6 | 3.32E+08 | 2.63E+08 | 1.78E+08 | 1.70E+08 | 1.67E+08 | 1.11E+07 | 6.48E+07 | 4.26E+07 | 2.16E+07 | 8.18E+05 |
| Replicate 7 | 3.34E+08 | 2.61E+08 | 1.63E+08 | 1.56E+08 | 1.52E+08 | 1.11E+07 | 6.53E+07 | 4.38E+07 | 1.63E+07 | 7.16E+05 |
| Replicate 8 | 3.44E+08 | 2.66E+08 | 1.88E+08 | 1.80E+08 | 1.73E+08 | 1.50E+07 | 7.27E+07 | 4.85E+07 | 2.26E+07 | 1.11E+06 |
| Replicate 9 | 4.14E+08 | 3.20E+08 | 2.07E+08 | 1.97E+08 | 1.90E+08 | 1.73E+07 | 8.03E+07 | 5.52E+07 | 2.41E+07 | 1.26E+06 |
| Replicate 10 | 5.36E+08 | 4.30E+08 | 2.20E+08 | 2.10E+08 | 1.98E+08 | 2.19E+07 | 1.12E+08 | 8.19E+07 | 1.42E+07 | 1.10E+06 |
| Replicate 11 | 2.33E+08 | 1.91E+08 | 1.14E+08 | 1.09E+08 | 1.05E+08 | 9.36E+06 | 5.97E+07 | 4.29E+07 | 9.85E+06 | 7.40E+05 |
| Pooled | 3.77E+09 | 2.93E+09 | 1.69E+09 | 1.61E+09 | 1.56E+09 | 1.33E+08 | 7.11E+08 | 4.87E+08 | 1.85E+08 | 9.66E+06 |

**Table S2.** Read statistics of the prometaphase wild-type samples generated in this study.

|  | Total | Mapped | Unique | Unique (HQ) | Cis | Trans heterolog | Cis 1kb+ | Cis 10kb+ | Cis sister | Trans sister |
| --- | --- | --- | --- | --- | --- | --- | --- | --- | --- | --- |
| Replicate 1 | 6.72E+07 | 5.28E+07 | 4.63E+07 | 4.37E+07 | 3.82E+07 | 8.01E+06 | 2.72E+07 | 2.42E+07 | 3.19E+06 | 2.99E+05 |
| Replicate 2 | 8.75E+07 | 6.45E+07 | 5.41E+07 | 5.10E+07 | 4.22E+07 | 1.19E+07 | 2.59E+07 | 2.26E+07 | 2.64E+06 | 2.23E+05 |
| Pooled | 1.55E+08 | 1.17E+08 | 1.00E+08 | 9.47E+07 | 8.05E+07 | 1.99E+07 | 5.31E+07 | 4.68E+07 | 5.84E+06 | 5.23E+05 |

**Table S3.** Read statistics of the NIPL-degraded samples generated in this study.

|  | Total | Mapped | Unique | Unique (HQ) | Cis | Trans heterolog | Cis 1kb+ | Cis 10kb+ | Cis sister | Trans sister |
| --- | --- | --- | --- | --- | --- | --- | --- | --- | --- | --- |
| Replicate 1 | 5.51E+08 | 4.27E+08 | 3.06E+08 | 2.93E+08 | 2.54E+08 | 5.24E+07 | 1.41E+08 | 9.78E+07 | 1.49E+07 | 8.86E+05 |
| Replicate 2 | 4.22E+08 | 3.35E+08 | 2.38E+08 | 2.27E+08 | 2.01E+08 | 3.65E+07 | 9.52E+07 | 6.47E+07 | 1.28E+07 | 9.73E+05 |
| Replicate 3 | 7.15E+07 | 5.19E+07 | 3.71E+07 | 3.53E+07 | 3.09E+07 | 6.20E+06 | 1.67E+07 | 1.20E+07 | 1.10E+06 | 1.09E+05 |
| Replicate 4 | 8.77E+07 | 6.54E+07 | 2.89E+07 | 2.73E+07 | 2.41E+07 | 4.81E+06 | 1.16E+07 | 8.32E+06 | 1.98E+06 | 2.16E+05 |
| Pooled | 1.13E+09 | 8.79E+08 | 6.10E+08 | 5.82E+08 | 5.10E+08 | 9.99E+07 | 2.65E+08 | 1.83E+08 | 3.07E+07 | 2.18E+06 |

**Table S4.** Read statistics of the Sororin-degraded samples generated in this study.

|  | Total | Mapped | Unique | Unique (HQ) | Cis | Trans heterolog | Cis 1kb+ | Cis 10kb+ | Cis sister | Trans sister |
| --- | --- | --- | --- | --- | --- | --- | --- | --- | --- | --- |
| Replicate 1 | 3.83E+08 | 3.10E+08 | 2.28E+08 | 2.19E+08 | 2.05E+08 | 2.36E+07 | 6.37E+07 | 4.49E+07 | 2.63E+07 | 8.86E+05 |
| Replicate 2 | 3.95E+08 | 3.21E+08 | 2.08E+08 | 1.98E+08 | 1.85E+08 | 2.28E+07 | 6.63E+07 | 4.73E+07 | 2.34E+07 | 9.73E+05 |
| Replicate 3 | 1.73E+08 | 1.39E+08 | 1.08E+08 | 1.03E+08 | 9.34E+07 | 1.49E+07 | 4.92E+07 | 3.61E+07 | 5.68E+06 | 3.44E+05 |
| Pooled | 9.51E+08 | 7.70E+08 | 5.44E+08 | 5.21E+08 | 4.83E+08 | 6.13E+07 | 1.79E+08 | 1.28E+08 | 5.54E+07 | 2.20E+06 |

**Table S5.**
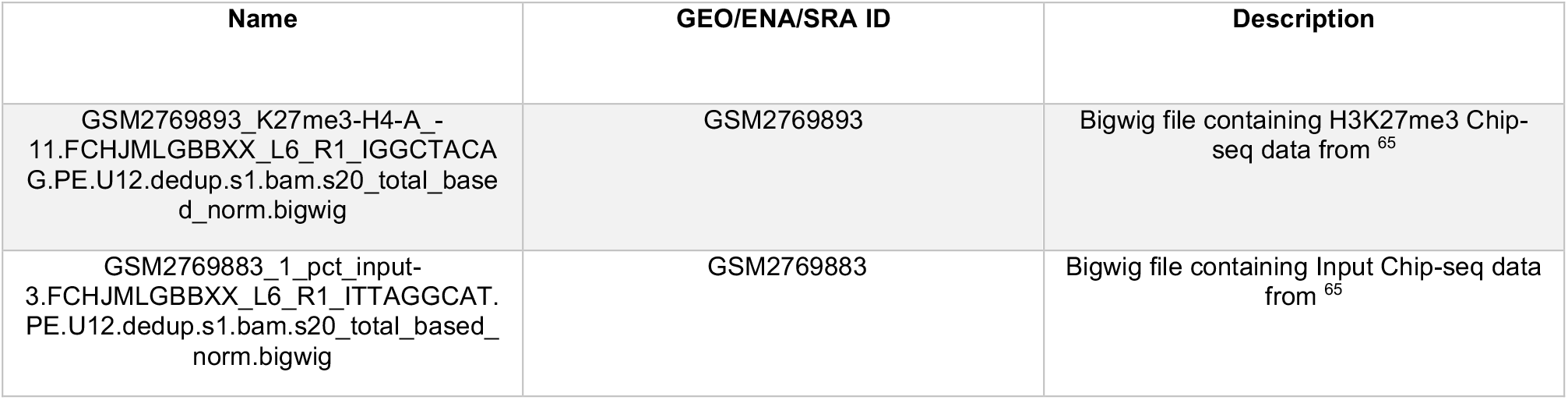
Published datasets used in this study.

**Table S6.** Cell lines used in this study.

| Name | Genotype | Plasmid used | Resistance marker |
| --- | --- | --- | --- |
| HeLa Kyoto | WT | - | - |
| HeLa Kyoto Sororin-AID | Tir1-3xMyc-T2A-Puro<br>(Lentiviral integration)<br>EGFP-AID-Sororin | Tir1-3xMyc-T2A-Puro-Lentivirus | Puromycin |
| HeLa Kyoto Nipbl-AID | Tir1-3xMyc-T2A-Puro<br>(Lentiviral integration)<br>AID-GFP-NIPBL<br>(Endogenously tagged) | Tir1-3xMyc-T2A-Puro-Lentivirus<br>AID-GFP-NIPBL-HR | Puromycin |
| HeLa Kyoto H2B-mCherry | H2B-mCherry<br>(Lentiviral integration) | Lenti-H2B-mCherry | Blasticidin |

**Table S7.** Plasmids used in this study.

| Name | Insert | Prokaryotic resistance marker | Eukaryotic resistance marker |
| --- | --- | --- | --- |
| pRRL | Tir1-3xMyc-T2A-Puro | Ampicillin | Puromycin |
| AID-GFP-NIPBL-HR | AID-GFP-NIPBL Repair template | Ampicillin | - |
| Lenti-H2B-mCherry | H2B-mCherry | Ampicillin | Blasticidin |
| EGFP-AID-Sororin-HR | EGFP-AID-Sororin Repair template | Ampicillin | - |

